# Snapshots of actin and tubulin folding inside the TRiC chaperonin

**DOI:** 10.1101/2021.03.26.436673

**Authors:** John J. Kelly, Dale Tranter, Els Pardon, Gamma Chi, Holger Kramer, Kelly M. Knee, Jay M. Janz, Jan Steyaert, Christine Bulawa, Ville O. Paavilainen, Juha T. Huiskonen, Wyatt W. Yue

## Abstract

The integrity of a cell’s proteome depends on correct folding of polypeptides by chaperonins. The TCP-1 ring chaperonin (TRiC) acts as obligate folder for >10% of cytosolic proteins, including cytoskeletal proteins actin and tubulin. While its architecture and how it recognises folding substrates is emerging from structural studies, the subsequent fate of substrates inside the TRiC chamber is not defined. We trapped endogenous human TRiC with substrates (actin, tubulin) and co-chaperone (PhLP2A) at different folding stages, for structure determination by cryogenic electron microscopy. The already-folded regions of client proteins are anchored at the chamber wall, positioning unstructured regions towards the central space to achieve their folding. Substrates engage with different sections of the chamber during the folding cycle, coupled to TRiC open-and-close transitions. Furthermore, the cochaperone PhLP2A modulates folding, acting as a molecular strut between substrate and TRiC chamber. Our structural snapshots piece together an emerging atomistic model of client protein folding through TRiC.

---

The group II chaperonin TRiC (also called chaperonin containing tailless complex, CCT) is essential for the folding and function of a growing list of proteins driving diverse cellular processes^1–4^. It is tasked to fold >10% of cytosolic proteins, including the essential actin and tubulin. TRiC also suppresses the misfolding and aggregation of neurotoxic proteins including huntingtin^5^ and α-synuclein^6^. The architecture of TRiC has been the subject of extensive x-ray crystallography and cryogenic electron microscopy (cryo-EM) studies^7–9^. Eight paralogous subunits (CCT1-CCT8) assemble into a hexadecamer of two back-to-back rings, enclosing a folding chamber^10^. This architecture facilitates nascent protein recognition at the apical domain with a built-in lid, as well as binding and hydrolysis of ATP in the equatorial and intermediate domains to drive lid closure for open-and-close conformational transition^11^. The evolutionary divergence to eight paralogous subunits further allows TRiC to fine-tune substrate specificity through differential recognition modes^12^ and rates of ATP binding and hydrolysis^13–15^.

Events that follow entrapment and confinement of nascent proteins in the chamber interior to assist folding into their functional states^10^, by contrast, remain sketchy, as atomic details of client protein interactions with TRiC interior are lacking. The two canonical TRiC substrates, actin and tubulin, are abundant filament-forming cytoskeletal proteins^16^. Due to their complex domain topology, these proteins depends on TRiC to achieve their native folds, and inherited mutations of actin and tubulin that disrupt TRiC engagement associate with human disease^17^. TRiC-mediated folding of actin and tubulin also requires the assistance of two cochaperone classes, prefoldin (PFD) and phosducin-like proteins (PhLPs). While PFD is tasked to stabilise nascent polypeptides emerging from ribosome for TRiC delivery^18^, PhLPs seem to modulate activity in the context of a TRiC-substrate-cochaperone ternary complex^19^. To obtain insights into folding of native substrates by TRiC, we set out to isolate endogenous human TRiC for cryo-EM structure determination. Our strategy aimed to entrap endogenous TRiC-bound substrates and cochaperones along the folding cycle, a feat not readily accomplished using reconstituted systems.

## Results

### Cryo-EM structure of endogenous TRiC complexed with subunit-specific nanobody

Using CRISPR/Cas9 knock-in, we inserted a C-terminal 3xFLAG into the *CCT5* genomic locus of human HEK293T cells, and affinity-purified endogenous TRiC (Extended Data Fig. 1). A ~50 kDa protein copurified with TRiC, which we identified as tubulin^20^ by liquid chromatography tandem mass spectrometry (LC-MS/MS) (Supplementary Data 1). Here, we prepared sample from the CCT5-FLAG cell line for solution digest LC-MS/MS, with wild-type un-transfected HEK293T cells as reference. We found that tubulin constitutes the most abundant peptides detected beyond TRiC subunits, followed by other substrate proteins and cochaperones (Supplementary Note 1).

To aid subunit alignment of pseudo-D8 symmetric TRiC during cryo-EM data processing, we raised a CCT5-specific nanobody (Nb18) by immunization of a llama with a recombinant homo-oligomeric form of CCT5^21^ (Extended Data Fig. 2). Nanobody Nb18 was verified to bind CCT5 *in vitro* by ELISA and bio-layer interferometry (BLI), and pulled down CCT5_16_ and TRiC from HEK293T by affinity column and co-eluted in size exclusion chromatography (Extended Data Fig. 2a-b). Nb18 did not bind other TRiC subunits when expressed recombinantly, such as CCT4 (Extended Data Fig. 2b, inset). Importantly, the binding of Nb18 did not interfere with either CCT5 or TRiC ATPase activity (Extended Data Fig. 2c).

Purified endogenous TRiC was complexed with ADP-AlF_x_ mimicking the post-hydrolysis state of the nucleotide. This potentially traps TRiC at the latter folding stages (*i.e*. prior to chamber opening and client release) where any bound substrates would have attained some structure. In addition, Nb18 was added before cryo-EM data collection (Supplementary Table 1). Image classification revealed 89% of particles (*i.e*. 2.5 million) in the closed state (closed-TRiC), resulting in a 2.5-Å resolution consensus map after imposing C2 symmetry (Fig. 1a, Extended Data Figs. 3,4). With this map, an atomic model of the entire TRiC–Nb18 complex was built (Fig. 1b). The eight paralogous subunits are assembled within each of the two rings as per previously reported arrangement (Fig. 1c)^22–24^ validated here by inter-subunit crosslinks (Supplementary Table 2), and by using Nb18 as a subunit-specific tag (Extended Data Fig. 2d, Supplementary Table 2).

**Fig. 1.**
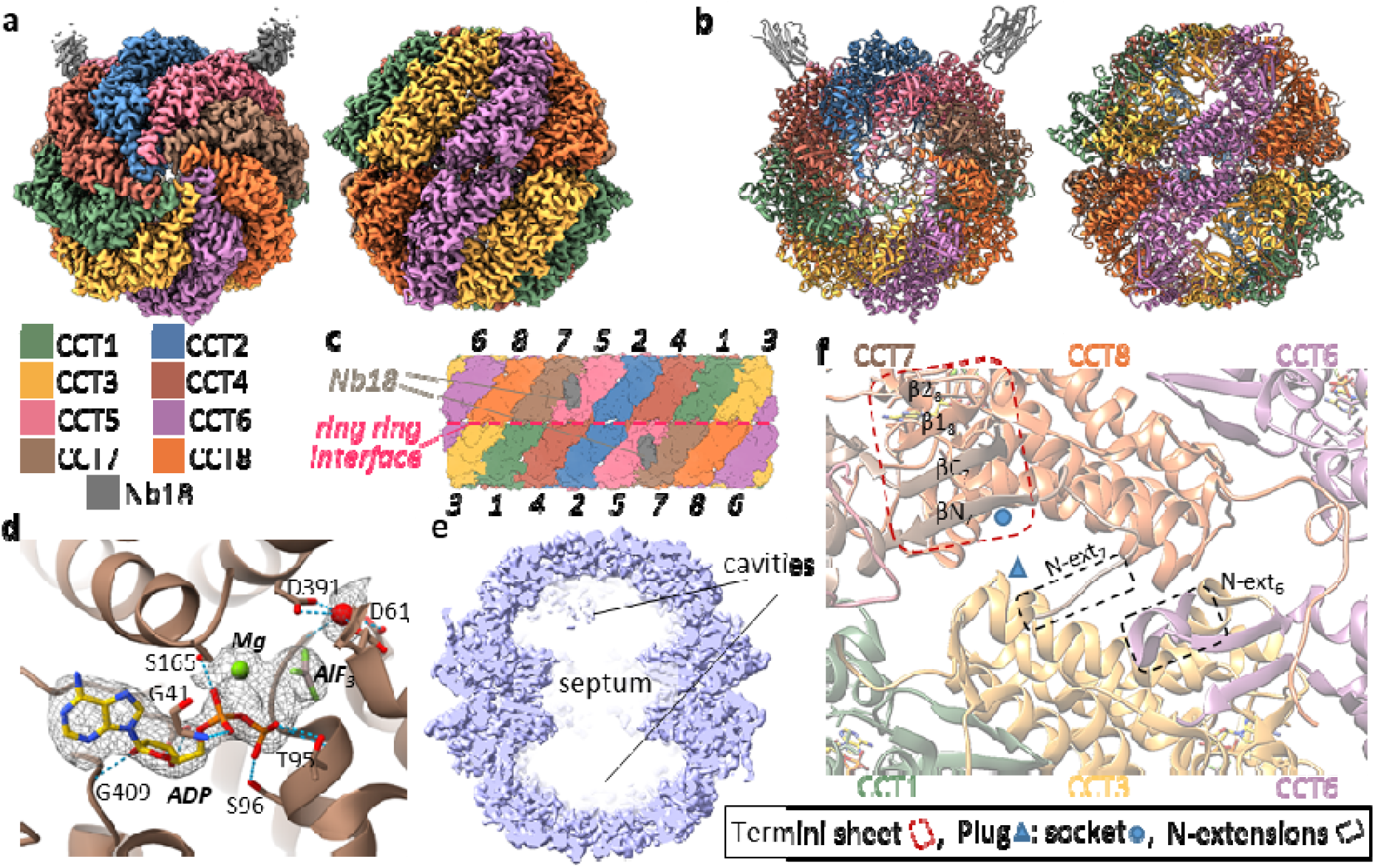
Structure of TRiC-Nb18 complex. (**a**) 2.5-Å resolution map of closed-TRiC in complex with Nb18 in top (*left*) and side (*right*) views. (**b**) Cartoon representation of closed-TRiC in same views as in panel *a*. (**c**) Flattened scheme of subunit arrangement for the two stacked rings. (**d**) Nucleotide binding site of CCT7 bound with ADP·Mg^2+^·AlF_3_. Conserved interacting residues are shown as sticks. (**e**) Side slice of closed-TRiC (fully empty class 4), highlighting the septum at the ring interface that creates two cavities in the TRiC interior. (**f**) Section of the ring interface, viewed from the interior, showing examples of intra-ring (termini-hairpin sheets) and inter-ring (plug-socket, N-extensions) contacts.

Nb18 is bound at only two locations of TRiC (one per copy of CCT5, Fig. 1a,b). The binding interface consists of CDR 1/2/3 loops of Nb18 and a hydrophobic patch at CCT5 equatorial domain, proximal (~15 Å) to the ATP binding site (Extended Data Fig. 2d). The Nb18-CCT5 interactions are mediated by Phe29, Arg53, and Trp101 from the CDR1, CDR2 and CDR3 loop regions, respectively (Extended Data Fig. 2e). The CCT5 epitope residues (e.g. Ile151, Met485, Pro487) are not conserved among other CCT subunits, suggesting that Nb18 binding is CCT5-specific. In agreement, Nb18 did not binding to recombinant CCT4 in BLI. CCT5-Nb18 crosslinks were also identified that confirmed the location of nanobody binding to CCT5 (Extended Data Fig. 2f). Our data also revealed no detectable presence of CCT5 homooligomers^21^ in the endogenous isolation from native HEK293 cells. Firstly, in all steps of TRiC isolation and purification, the proportion of FLAG-tagged CCT5 remained stoichiometric with other CCT subunits. Secondly, cryo-EM 2D and 3D classifications did not present any evidence of CCT5 homo-oligomeric complexes bound with more than two nanobodies per hexadecamer.

The classical TRiC features^25,26^ are illustrated here in atomic details (Extended Data Fig. 6a), including conserved domain motifs, and ATP-binding sites that are occupied here by Mg^2+^·ADP·AlF_x_ and a water molecule (Fig. 1d, Extended Data Fig. 6b). Importantly, our closed-TRiC model provides unprecedented clarity to inter-subunit contacts within ring (*cis*) and across rings (*trans*) (Fig. 1f), through three salient features. First, intra-ring *cis* contacts are mediated by the N-/C-terminal β-strand of one subunit with β1-β2 hairpin from the adjacent subunit^27,28^, forming concerted 4-stranded β-sheets within each of the two rings. The intra-ring sheet formation is fully visualised in our model and the consequence is a rigid septum mid-level between rings, constricting the interior into two largely separate cavities (one per ring)(Fig. 1e). Second, inter-ring *trans* interactions are largely maintained at the ring-ring interface by fitting the ‘plug’ (α4-α5 linker) of one subunit, into the ‘socket’ (α14-linker-α15) of the subunit *in trans* that it stacks with (Fig. 1f, Extended Data Fig. 7c). Third, our model unravels an ordered-coil preceding the N-terminal β-strand of each subunit, reaching out to the equivalent coil of not its *trans* stacked subunit, but to the +2 subunit in the anticlockwise direction (Fig. 1f, Extended Data Fig. 7b, top). The exception to this N-terminal inter-ring network is CCT4/CCT4’, where its N-terminus points to the solvent exterior.

### Tubulin-bound TRiC in the closed state

The closed-TRiC consensus map revealed additional density within the folding cavity (Fig. 1e, Extended Data Fig. 4), potentially representing a mix of native substrates. To resolve these additional densities, we performed another 3D classification on the closed particles, and simultaneously relaxed two-fold symmetry^29^. Particles in the most abundant class (29.7% total closed particles; Extended Data Fig. 3) were refined to generate a 3.0-Å reconstruction, where we identified the additional density as tubulin (Fig. 2a, left). In our LC-MS/MS analysis, many isoforms of tubulin were highly enriched in the CCT5-FLAG cell line, and six of the top 25 most enriched peptides included different forms of β-tubulin (Supplementary Data 1, Extended Data Fig. 5). Thus, the tubulin density likely reflects a mixture of different tubulin forms including α-, β-, and γ-tubulin (sharing 31-41% sequence identity), although several isoforms of β-tubulin were among the most enriched peptides from affinity purification. We therefore built our model based on tubulin beta-2A chain (TUBB2A), the highest enriched tubulin peptide.

**Fig. 2.**
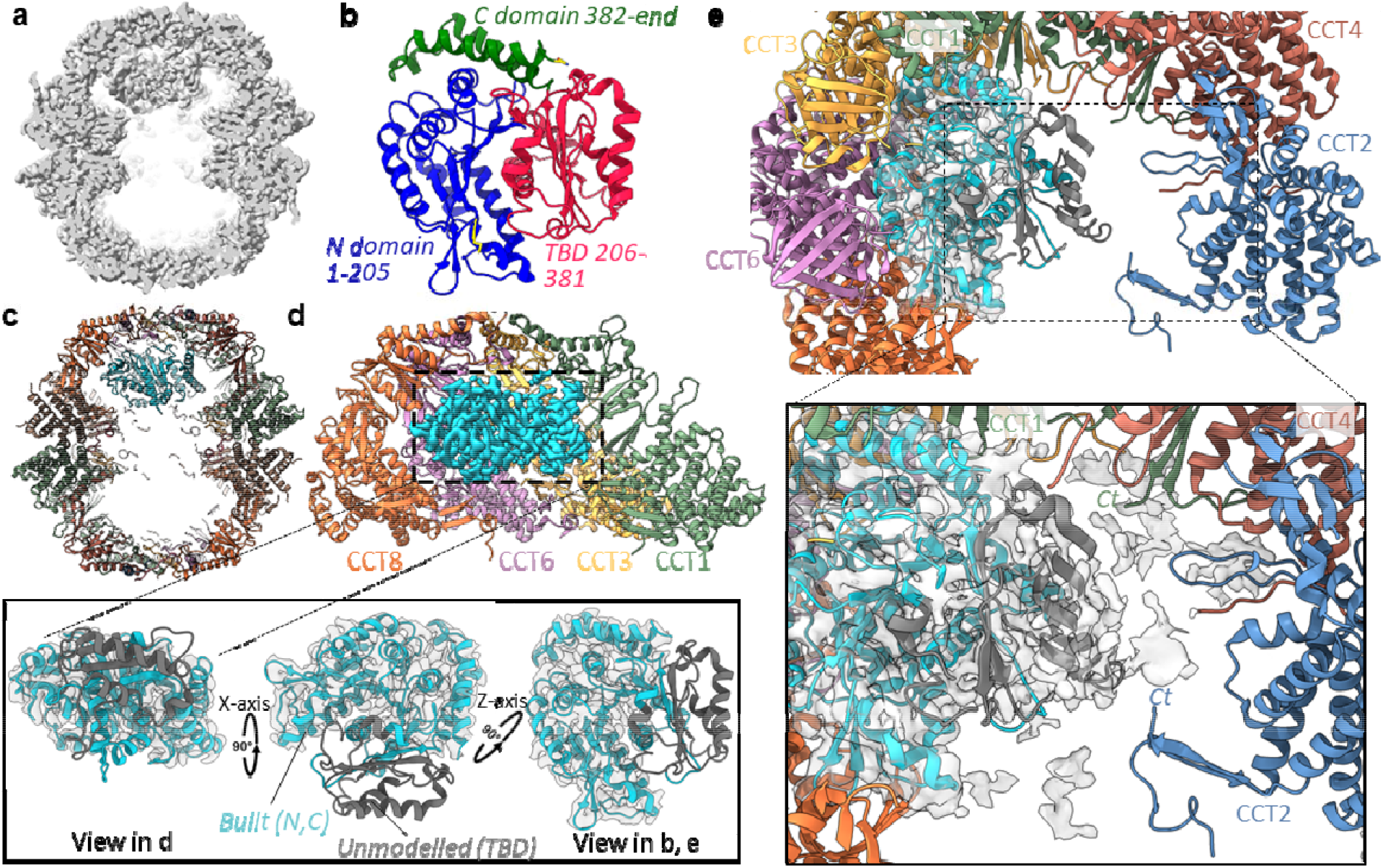
TRiC-tubulin complex. (**a**) Side slice map for closed-TRiC showing substrate density within one cavity of chamber. (**b**) Topology of native β-tubulin (PDB: 6I2I). (**c**) Overall model of TRiC-tubulin complex. (**d**) Slice of the CCT8-CCT6-CCT3-CCT1 hemisphere from one ring, in contact with the substrate density (cyan). *Inset*: Three orthogonal views showing region of β-tubulin built into the substrate density (cyan). Unmodelled TBD from tubulin (orange) is shown for reference. (**e**) Top-down view of chamber interior, with tubulin density (cyan) adjacent to one hemisphere, and space in the interior that can accommodate the unmodelled TBD. *Inset*: At lower isosurface threshold, substrate density (grey surface) can accommodate the TBD (dark grey, taken from PDB 6I2I) which extends towards the C-termini of CCT1 and CCT2.

A tubulin monomer comprises the N-terminal nucleotide-binding domain, taxol-binding domain (TBD), and C-terminal domain (Fig. 2b)^30^. We traced the N- and C-terminal domains of β-tubulin, which adopt a nearly folded conformation (Fig. 2d, inset), while the TBD is largely disordered. Tubulin is localised at the inner walls of one cavity, to the level of apical domains (Fig. 2c,d). The N- and C-terminal domains interact mainly with CCT3 and CCT6, and partly with CCT8 and CCT1 (Fig. 2d,e). These interactions are mediated by the helical protrusions and few loops in the apical domain that line the inner wall, with minor interactions involving intermediate and equatorial domains (Supplementary Note 2, Extended Data Fig. 8). Of note, CCT3/6/8 subunits constitute the hemisphere identified with low ATP binding and hydrolysis rates^14^. At lower isosurface threshold, the largely disordered TBD is seen to extend into the cavity centre (Fig. 2e, inset), towards the other hemisphere (CCT4/2/5/7) that exhibits strong ATP binding and hydrolysis, with potential interactions with C-termini of CCT1 and CCT2. Altogether, for the first time, a tubulin folding intermediate is observed in atomic details inside the TRiC chamber, where nearly folded regions are held by interactions with the TRiC inner walls, facilitating unfolded regions to attain native structure in the central space.

### A TRiC–actin–cochaperone complex

Intriguingly, another symmetry-relaxed 3D class (20.0% of all closed particles; Extended Data Fig. 3) presented density within both ring cavities (Fig. 3a), clearly different from the first class with tubulin. Another canonical substrate of TRiC, actin, could account for part of the density, into which we indeed built an actin monomer based on its detection in LC-MS/MS (Supplementary Data 1). Though actin was not enriched to the degree of tubulin and PhLP2A in LC-MS/MS, actin was slightly enriched in two peptide identifications including β-actin (ACTB) and γ-actin (ACTG1) (Extended Data Fig. 5b). Side chain analysis determined that cytoplasmic actin 1 (β-actin, ACTB), and 2 (γ-actin, ACTG1) were the most likely isoforms present in the sample. ACTB and ACTG1 only differ in their first ten residues, most of which are disordered in our model. We therefore used ACTB to build our atomic model, though the sample is likely to be a mixture between both isoforms of cytoplasmic actin.

**Fig. 3.**
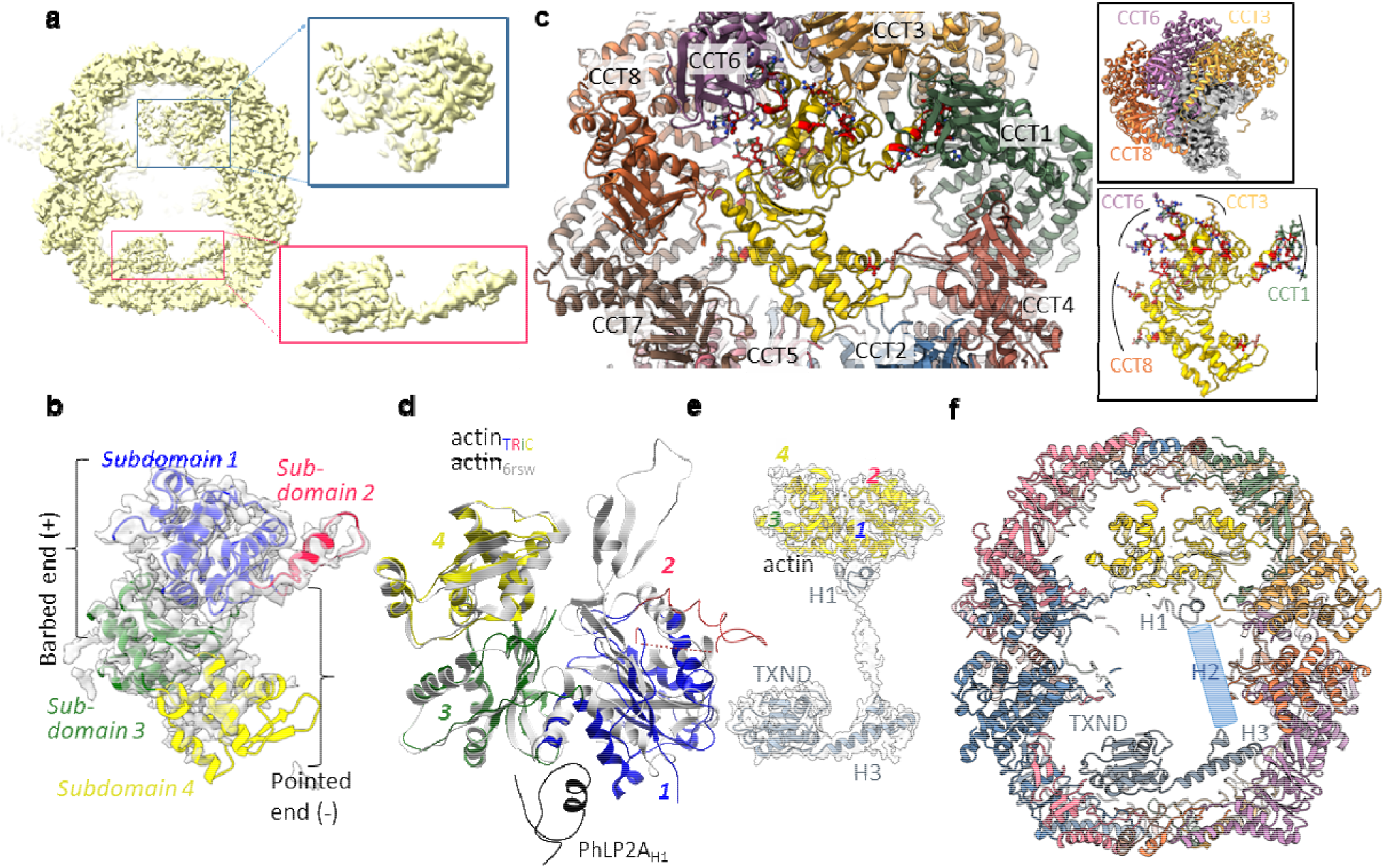
TRiC-actin-cochaperone complex. (**a)** Side slice map of closed-TRiC showing protein density in both chamber cavities. (**b**) Top-down view of actin model modelled from map density (grey), coloured by subdomains. (**c**) Top-down view of chamber interior showing TRiC subunits in contact with actin (yellow). *Inset*, *top*: Three subunits form main contacts with actin (grey). *Inset*, *bottom*: Subunit side-chains (sticks) in contact with actin (yellow). (**d**) Overlay of our actin model with native actin structure (PDB 6RSW). (**e**) Non-TRiC map density at lower threshold that can accommodate actin (yellow) bound with PhLP2A (grey). (**f**) Overall model of TRiC-actin-PhLP2A ternary complex. PhLP2A helix H2 is not modelled and shown as a cartoon cylinder.

Native actin monomer folds into subdomains 1-4 with an inter-domain ATP-binding cleft. Our model of actin, positioned similarly to tubulin (Fig. 2a, 3a), depicts again a partly folded state. Subdomains 1 and 3, known as the ‘barbed end’, are well defined. Subdomain 1 is localised close to CCT3 and CCT6, while subdomain 3 contacts extensively the hairpin-termini of CCT2, CCT7 and CCT8 (Fig. 3c, Extended Data Fig. 9a,b,c,g, Supplementary Note 3). By contrast, subdomains 2 and 4, known as the ‘pointed end’, are more disordered (Fig. 3b), making less contact to TRiC than the barbed end (Fig. 3c and bottom inset). No density is observed for the subdomain 2 D-loop, a region undergoing disorder-to-order transition during actin polymerisation. D-loop, as it attains tertiary structure, can extend into a groove formed in the CCT1 intermediate domain (Extended Data Fig. 9d,e). Observed density for subdomain 4 is weak (Fig. 3b), from which we built a C-α trace that reveals minimal TRiC contact with CCT4 apical domain (Fig. 3c, Extended Data Fig. 9f).

Additional density remained beneath actin subdomains 1 and 3 (Fig. 3e). At low isosurface threshold, this is linked up, by cylindrical density spanning the inter-ring septum, to the density in the other, *trans* cavity (Fig. 3a,e). We hypothesised that these connected densities could represent an actin-binding partner e.g. one of the phosducin-like proteins (PhLPs). Among the five human PhLPs, PhLP2A and the homologous PhLP2B have been shown to regulate actin and tubulin folding^31^; importantly, PhLP2A was enriched in our LC-MS/MS analysis (Supplementary Data 1, Extended Data Fig. 5). The PhLP2A sequence comprises an N-terminal helical domain with three predicted helices H1-H3 and a C-terminal thioredoxin domain (TXND) (Extended Data Fig. 10a). We unambiguously identified PhLP2A helix H1, helix H3 and its preceding loop, in addition to TXND, and built their atomic models into the density (Fig. 3e).

The PhLP2A N-terminal helix H1, the most sequence-divergent region among PhLPs^19^, fits into the *cis*-ring density proximal to actin, anchoring the subdomain 1-3 interface (Fig. 3f, Extended Data Fig. 10b). Consequently, actin subdomain 1 has rotated by nearly 40° away from the core, relative to native monomeric actin (Fig. 3d). Helix H1 in the *cis*-ring is connected to the bulk of PhLP2A in the *trans*-ring by weak cylindrical density that can accommodate helix H2. While not modelled in our structure, helix H2 would traverse >60 Å longitudinally from the *cis*-ring through the septum to the *trans*-cavity, making minimal TRiC contacts (Fig. 3f, Extended Data Fig. 10c). Upon reaching the level of the CCT2’ and CCT3’ apical domains in the *trans* cavity, helix H2 is followed by a loop region and helix H3 which traverses 35 Å latitudinally to the other hemisphere of this cavity close to CCT6, reaching the C-terminal TXND (Fig. 3f, Extended Data Fig. 10d). This TXND fold, highly conserved across all PhLPs and superimposable with the PhLP2B TXND structure^32^, contacts TRiC through the CCT5 apical domain loops, and partly the flanking subunits of CCT2, CCT7 and CCT4 (Fig. 3f). The PhLP2A 22-aa C-terminus was disordered, but proximal to the termini extension of CCT3, CCT1 and CCT4. Altogether, this ternary complex structure confirms PhLP2A as a binding partner of actin, and depicts PhLP2A as a molecular ‘strut’, anchored to the TRiC *trans* cavity, while reaching out to hold and stabilise the actin folding intermediate in the *cis* cavity.

### Substrates bind differently to TRiC in the open state

Beyond the closed states, initial 3D classification revealed TRiC in the open state (open-TRiC) for 10.7% of particles (Extended Data Fig. 3). These generated a consensus map at 4.0-Å resolution (Fig. 4a), ranging locally from 3.5 Å (equatorial domains) to 9.0 Å (apical domains) (Extended Data Fig. 4). We built into the ordered map portion all equatorial domains up to the ATP binding pockets, which are again bound with Mg^2+^·ADP·AlF_x_ except CCT4 and CCT5, where the smaller ligand density is consistent with bound ADP (Extended Data Fig. 6c). We then modelled apical domains to the less ordered map portion by molecular dynamics flexible fitting. This open-TRiC conformation (Fig. 4b), previously seen with AMPPNP-bound yeast CCT, substrate-bound bovine CCT, and substrate mLST8-bound human CCT^23,33^, is the consequence of concerted rigid-body rearrangement of all (apical, intermediate, equatorial) domains, relative to closed-TRiC, to varying degrees among different subunits (Extended Data Fig. 7a). Such open-and-closed transitions impart major consequences on inter-subunit as well as TRiC-substrate contacts.

**Fig. 4.**
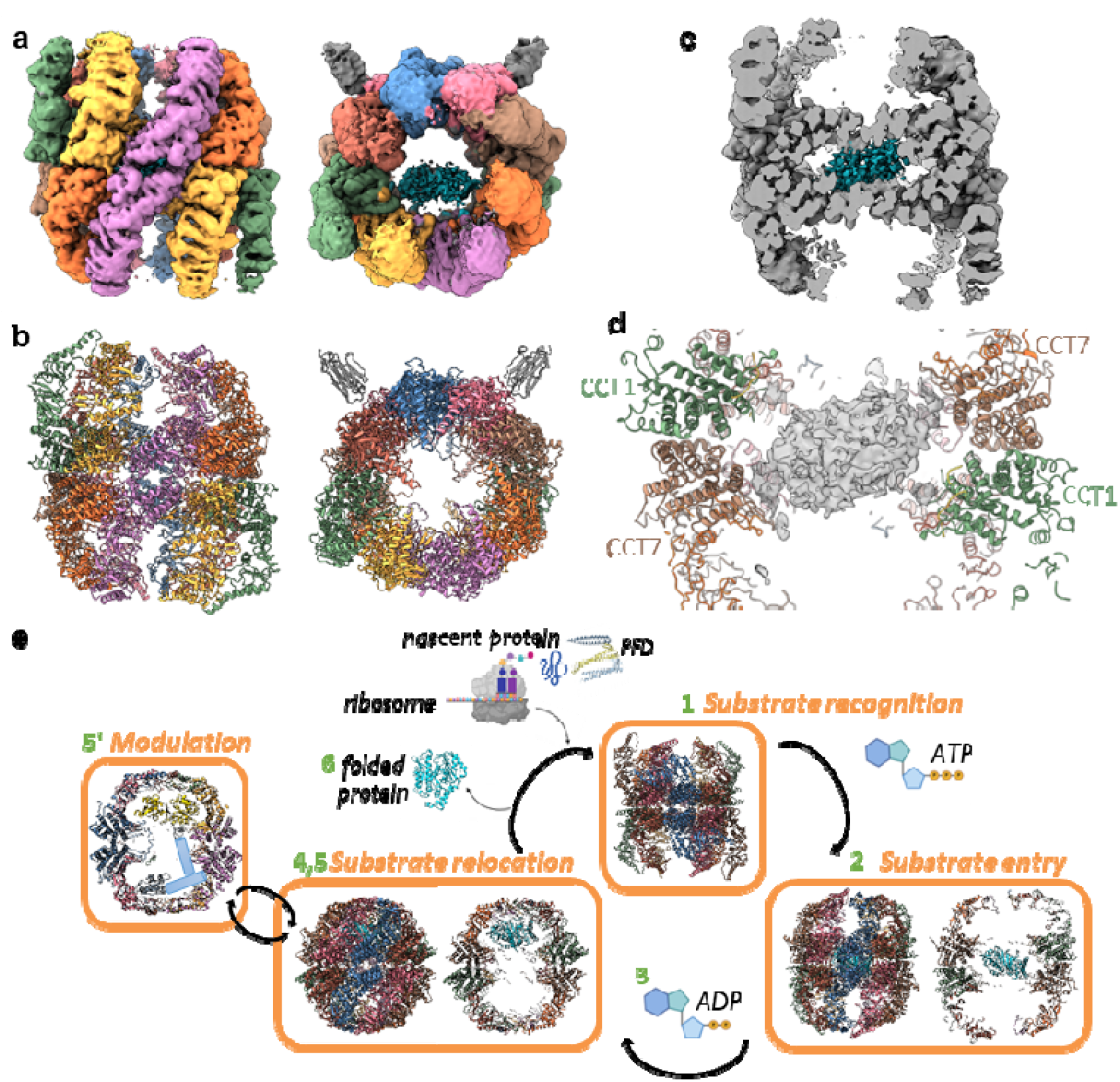
Substrate density in open-TRiC. (**a**) 4.0-Å resolution reconstruction map of open-TRiC in side (*left*) and top (*right*) views. (**b**) Cartoon representation showing apical domains built into map, as well as intermediate and apical domains modelled by flexible fitting; same views as panel *a*. (**c**) Side slice of open-TRiC with substrate density in cyan. (**d**) Slice of substrate density at low isosurface threshold, in the septum of open-TRiC. (**e**) Proposed mechanism of TRiC-mediated substrate folding based on this work. Numbers refer to steps described in text. PFD, prefoldin.

In open-TRiC, a significant portion of the N-terminus is disordered in most subunits (Extended Data Fig. 7b, bottom). Therefore, unlike closed-TRiC, the pre-strand coil is not available for inter-ring contacts, and the N-terminal β-strand is missing from intra-ring sheet formation (which is now 3-stranded). Additionally, the inter-ring plug-socket interactions seen in closed-TRiC (Extended Data Fig. 7c) have now disengaged in open-TRiC with a rearranged interface (Extended Data Fig. 7d).

The rearranged intra-/inter-subunit contacts in open-TRiC indicate that the septum structure at the ring-ring interface has become more dynamic, and less constricted between the two cavities, when compared to closed-TRiC (Fig. 4c). Indeed, our open-TRiC map revealed weak substrate density at the level of the equatorial domains, surrounded by the less rigid septum (Fig. 4d). This density is positioned >30 Å deeper into the chamber when compared to the actin and tubulin positions in closed-TRiC, occupying the same site as mLST8 substrate in the human open-TRiC structure^33^. We reason that this density represents a summed average of different substrates sampled in the data set, and cannot be attributed to a single substrate. This density is in close proximity with CCT7 subunits from both rings (Fig. 4d), while engaging more transiently the dynamic termini/septum of other subunits.

## Discussion

Our endogenous isolation strategy has captured TRiC-bound actin and tubulin, abundant proteins in the cell, during their acts of folding, while the subunit-specific nanobody and improved 3D image classification have enhanced the resolution beyond recent TRiC structures^18,26,33^. Our data illuminate a mechanism whereby a substrate traverses different sections of TRiC, dependent on its folding status, and in coordination with an ATP-driven conformational cascade within the chamber (Fig. 4e).

Previous studies have demonstrated that nascent actin and tubulin polypeptides are recognised at the apical domain of nucleotide-free TRiC^12,34^ and released into the chamber by the ATP-induced lid formation^35^. Here, our novel snapshots map out the subsequent events inside the chamber. Our open-TRiC model suggests that partly-folded substrates, once inside the chamber, are sequestered and loosely held in the septum. This agrees with other structural studies of open-TRiC^33^, and recent work showing that actin engages TRiC having already adopted some degree of structures^34^. Therefore, the septum could serve as the reception point upon substrate entry into the chamber.

Our closed-TRiC models in the post-hydrolysis state reveal the result of long-range conformational changes that occur to TRiC upon ATP-hydrolysis, which extends from lid closure at the apical domains all the way to structural rearrangement of the septum in the central chamber. Consequently, a combination of near-native structure and its steric hindrance at the rearranged septum has translocated and confined the folding substrate into one of the two chamber cavities. Here, subunit-specific contacts stabilise the structured regions of substrates ^34^ (e.g. actin subdomains 1&3, tubulin N-/C-domains), allowing the less folded regions to achieve native structure, presumably by utilising chamber space and additional subunit contacts if necessary.

Furthermore, our TRiC-actin-PhLP2A model reveals for the first time the structure of full-length PhLP2A, and shows that this cochaperone plays two roles in TRiC quality control, via extensive helical segments. By holding the substrate within closed-TRiC (as a *strut*) until a folded state is reached for chamber release (as a *sensor*), the cochaperone therefore prevents pre-mature chamber opening and abortive folding. This provides a mechanistic explanation on the role of phosducin-like cochaperones in modulating the rate of ATP hydrolysis that governs TRiC open-and-close transition^31^. Altogether, this work sets the stage for future characterisation of other TRiC substrates (e.g. WD40 proteins, β-propellers) and cochaperones (e.g. PFD) to piece together the inner workings of this nanomachine.

## METHODS

No statistical methods were used to predetermine sample size. The experiments were not randomized. The investigators were not blinded to allocation during experiments and outcome assessment.

### Tagging of endogenous TRiC/CCT with CRISPR/Cas9

We chose to target CCT5 for purification tag insertion to the endogenous TRiC/CCT. To insert a purification tag to the endogenous TRiC/CCT, we transfected HEK293T cells with Cas9 and sgRNA targeting the gene encoding for the CCT5 subunit (Addgene eSpCas9(1.1) vector) along with a double stranded DNA fragment for a donor template containing a 3xFLAG/SpyTag^36^ to be inserted at the C-terminus of CCT5. We chose to target CCT5 due to previous success in expression and purification of recombinant CCT5 that is His-tagged at the C-terminus^21^. Insertion of a tag to the endogenous TRiC/CCT was achieved by a marker-free coselection strategy^37^. HEK293T cells were transfected with plasmid encoding eSpCas9(1.1) and sgRNAs targeting *CCT5* and *ATP1A1* (modified from Addgene #86613) along with linear dsDNAs to act as HDR templates by which to insert 3xFLAG/Spytag at the C-terminus of *CCT5*, as well as the Q118R and N129D ouabain resistance conferring point mutations to *ATP1A1*. Following ouabain selection, the polyclonal cell pool was assessed for FLAG-tagged CCT5 via immunoblotting. Single cells from confirmed pools were seeded into 96-well plates by FACS and resultant monoclonal lines were again screened via in-cell Western assay. Three positive monoclonal lines were identified and each was further verified by PCR, sequencing, and Western Blots.

### Llama immunization, nanobody selection, and nanobody purification

We raised CCT5-specific nanobodies by immunization of a llama^38^, using as antigen the recombinant *E. coli*-expressed human CCT5, which was previously shown to exist as a TRiC-like homo-hexadecamer (CCT5_16_)^21^. The llama received 6 injections of the antigen over a span of 6 weeks, after which point the ORF of nanobodies were isolated by extracting blood, separating out peripheral blood lymphocytes, and isolating RNA for use in creating cDNA libraries. *In vitro* selection of nanobodies was performed using phage display and binders were evaluated by enzyme-linked immunosorbent assay (ELISA) detecting presence of nanobody in the sample. 28 nanobodies from 18 families were identified, among which nanobody Nb18 was selected for cryo-EM experiments based on characterisation by pull-down and biolayer interferometry studies (Extended Data Fig. 2). Nb18 (ref no. CA14679) is constructed in a pMESy4 vector incorporating C-terminal His6 and CaptureSelect C-tag (or EPEA-tag), which allows recombinant expression in *E. coli*, and purification by Ni-NTA affinity and size-exclusion chromatography.

### Isolation and purification of human TRiC

Monoclonal HEK293T cells confirmed to contain FLAG-tagged CCT5 were grown in suspension and harvested by freeze-thaw of cell pellet and resuspended in lysis buffer (100 mM HEPES pH 7.5, 100 mM NaCl, protease inhibitors, 1 mM DTT, benzonase, and 1% Triton X-100) and incubated at 4°C for one hour. Lysate was centrifuged at 35,000 x g and supernatant was passed through a 0.4 μm filter and was incubated at 4°C for one hour with 5ml of anti-FLAG resin (Anti-DYKDDDK Tag (L5) Affinity Gel Protocol BioLegend cat no. 651503). Resin/lysate mixture was passed through a gravity column, washed with 100CV of wash buffer (20 mM HEPES pH 7.5, 100 mM NaCl) and eluted with wash buffer containing 0.15 mg/ml 3x FLAG peptide (Sigma cat no. F4799) (Extended Data Fig 1b). The elutates were concentrated and ran through a Superose 6 10/300 size exclusion chromatography column (Extended Data Fig 1c-d). Fractions containing purified TRiC were concentrated and flash frozen and stored at −80°C.

### Cryo-EM structure determination

Purified TRiC/CCT complex (1.5 mg/ml) was premixed with Nb18 at a 1:1 molar ratio, with 1 mM ATP, 5 mM MgCl_2_ and AlF_x_ (5 mM Al(NO_3_)_3_ and 30 mM NaF^11^. A 3-μl aliquot was pipetted on a holey carbon cryo-EM grid (1.3/1.2 Cu 200 mesh, Quantifoil) and vitrified by plunge-freezing in liquid ethane (Vitrobot Mark IV, Thermo Fisher Scientific). The data was collected using a 300-kV transmission electron microscope (Titan Krios, TFS) on a direct electron detector (Gatan K3). Data collection parameters are in Supplementary Table 1.

Cryo-EM data were processed in Scipion software framework^39^. Movie frames were aligned, dose weighted and averaged using MotionCor2^40^. Contrast transfer function (CTF) parameters were estimated using CTFfind4^41^. Particles were picked using crYOLO and subjected to several rounds of classification in RELION^42–44^. The particles were classified to closed and open states first using 2D classification. The initial 3D model for each state was calculated *ab initio*. Particles were further 3D classified in four separate sets (Extended Data Fig. 3). The closed particles were further classified to different substratebound states using option –relax_sym C2 in RELION^45^. This revealed actin–co-chaperone class which was not captured using conventional 3D classification in RELION. Particles in the selected classes, including the consensus map combining all classes with C2 symmetry imposed, were refined using the gold-standard refinement protocol in RELION. The resolution of the final cryo-EM maps was estimated by Fourier shell correlation (FSC) combined with phase randomization to account for the effects of masking. Data processing parameters are in Supplementary Table 1.

### Model building, refinement and validation

To model the atomic structure of TRiC/CCT closed state, a homology model was created for each chain in SWISS-MODEL using the yeast structure (PDB: 6KS6) as a template. A model for Nb18 was created in a similar manner (PDB: 6QX4 chain C). These models were fitted to the 2.5-Å resolution cryo-EM map of the closed state maximising real space cross-correlation in UCSF Chimera. The tubulin (PDB:6I2I chain B) and actin (PDB:2Q1N) structures were fit in a similar way to the 3.0-Å resolution closed map. Models were manually adjusted in COOT and subjected to several rounds of real space refinement in PHENIX. To model the open state, the previously solved open state (PDB:6NRA) model was first fitted as a rigid body to the 3.5-Å resolution cryo-EM map of the open state and then manually built in COOT. Molecular dynamics flexible fitting was performed using Flex-EM software^46,47^ to create an open model containing apical domains that roughly fit the 4.0-Å reconstruction of the open state. Model refinement and validation statistics are in Supplementary Table 1.

Tryptic digest mass spectrometry (MSMS) and side chain modelling confirmed that the predominant copy of CCT6 in the TRiC complex isolated from HEK293T is the ubiquitously expressed CCT6A isoform, and not the alternatively spliced isoform CCT6B^48^. The predominant peptide coverage in the MSMS data was for CCT6A, however CCT6B was detected as the 22^nd^ most abundant protein. We did not observe any density for the affinity tag inserted at the C-terminus of CCT5, although we demonstrated fully intact FLAG-tag identified by western blot, SDS-PAGE analysis, and FLAG-affinity mediated pull-down of the TRiC complex (Extended Data Fig 1a-c).

### Binding and activity assays

Biolayer inteferometry (BLI) experiments were performed on a 16-channel ForteBio Octet RED384 instrument at 25 °C, in buffer containing 20 mM HEPES pH 7.5, 150 mM NaCl, and 0.1% BSA. 50 μL of 100 ng/uL biotinylated CCT5 was loaded to the streptavin coated sensors. The concentration for Nb18 used ranged from 40 μM to 39 nM. Measurements were performed using a 300 second association step followed by a 300 second dissociation step on a black 384-well plate with tilted bottom (ForteBio) using a serial dip method from low to high Nb concentration. The baseline was stabilized for 30 sec prior to association and signal from the reference sensors was subtracted and steady-state kinetics were fit using Octet Data Analysis software.

Malachite green ATPase activity assays were performed by incubating protein samples in ATPase assay buffer (20 mM HEPES pH 7.5, 150 mM KCl, 5 mM MgCl2). TRiC samples were diluted to 0.5 μM and incubated for 30 minutes in the presence of 250 μM ATP in a final volume of 50 μL. 100 μL of malachite green reagent was added to the sample wells and incubated for an additional 30 minutes. Absorbance at 620 nm was read and the amount of free phosphate released (ρmol) was quantified using a phosphate standard curve. Data was reported using average and standard deviation of discrete sample replicates.

### Liquid chromatography with tandem mass spectrometry

Quadruplicate samples of WT HEK293T and CCT5-FLAG tagged cells were lysed and purified using anti-FLAG affinity matrix (Biolegend cat no. 651501) using 20mM HEPES pH 7.5 and 100mM NaCl wash buffer and 200mM Glycine/150mM NaCl, pH 2.2 elution buffer. AP-MS elutions were reduced in volume in a speedvac (2.5 hours, 56C) and re-suspended in 50 μL 7.2 M urea, 100 mM NH4HCO3 and incubated for 15 min at 25°C. Cysteines were reduced with 10 mM DTT (0.5 uL of 1M stock) for 1 hr at 51°C and then protected with 30 mM iodoacetamide (Sigma, I1149-25G; 4.2 uL of 0.36M stock) for 45 min in the dark at 25°C. The reaction was quenched with 25 mM DTT and urea was diluted to < 1 M with 50 mM NH_4_HCO_3_. Added MS-grade trypsin (Thermo, MS grade) in a ratio of 1:20 (trypsin:protein, w:w) to solution and incubated for 16 hr overnight at 37°C. Trypsinized peptides were desalted using C18 desalting pipette tips (ThermoFisher, 87782). Peptides were injected (2.0 μL) and separated using an Ultimate 3000 RSLC nano liquid chromatography system (Thermo Scientific) coupled to a LTQ Orbitrap Velos mass spectrometer (Thermo Scientific) via an EASY-Spray source. Sample volumes were loaded onto a trap column (Thermo Scientific, cat no. 164564) at 8 ul/min in 2% acetonitrile, 0.1% TFA). Peptides were eluted on-line to a 50cm analytical column (Thermo Scientific, cat no. ES803). Separations were carried out using a ramped 120 min gradient from 1-90% buffer B (buffer A: 5% DMSO, 0.1% formic acid; buffer B: 75% acetonitrile, 5% DMSO, 0.1% formic acid). The mass spectrometer was operated in positive polarity using a data-dependent acquisition mode. Ions for fragmentation were determined from an initial MS1 survey scan at 30,000 resolution (at m/z 200) followed by Collision-induced dissociation (CID) of the top 10 most abundant ions with a normalised collision energy of 35. A survey scan m/z range of 350 – 1500 m/z was used and charge state exclusion enabled for unassigned, +1, +8 and >+8 ions. Lock mass correction was enabled using the following ions: 401.92272 and 445.12003.

Data was processed using the MaxQuant^49,50^ software platform (v1.6.2.3) with database searches carried out by the in-built Andromeda search engine against the Uniprot homo sapiens database (v20180104; 161,549 entries). A reverse decoy database was used at a 1% false discovery rate (FDR) for peptide spectrum matches and protein identification. Data was visualised and analysed further in Perseus^51^ (version 1.6.2.2).

For XL-MS samples experiment was repeated as above after the following initial steps: A 5 mg: 5 mg aliquot of isotopically-coded BS3 d0/d4crosslinker (Thermo) was reconstituted to 25 mM in water and immediately added to 25ug TRiC in the optimal ratio determined experimentally (between equimolar amounts to the number of moles of lysine residues to 10X the number of lysines). The crosslinking reaction was incubated for 30 min at 25°C with mild agitation and the reaction was quenched with 50 mM NH4HCO3 (1:20 dilution from 1M stock, or 5 uL) for 20 min at 25°C.

## End Notes

## Acknowledgements

The Structural Genomics Consortium is a registered charity (Number 1097737) that receives funds from AbbVie, Bayer Pharma AG, Boehringer Ingelheim, Canada Foundation for Innovation, Eshelman Institute for Innovation, Genome Canada, Innovative Medicines Initiative (EU/EFPIA) [ULTRA-DD grant no. 115766], Janssen, Merck & Co., Novartis Pharma AG, Ontario Ministry of Economic Development and Innovation, Pfizer, São Paulo Research Foundation-FAPESP, Takeda, and Wellcome Trust [092809/Z/10/Z]. J.J.K. is supported by the Nuffield Department of Medicine DPhil Prize Studentship. The work by J.J.K. and W.W.Y. is also supported by funding from Pfizer Inc. This project was also funded by the Academy of Finland consortium grant SEMMA (314669 to J.T.H, 314672 to V.O.P.). Cryo-EM was carried out with support of the Biocenter Finland and Instruct-HiLIFE cryo-EM core facility (University of Helsinki), Oxford Particle Imaging Centre (University of Oxford) and eBIC (Diamond Light Source, UK, visits EM20223-30 and EM20223-31). We thank Benita Löflund and Pasi Laurinmäki (University of Helsinki), Beth MacLean (University of Oxford), Julika Radecke and Andy Howe (eBIC) for technical assistance with cryo-EM experiments, and CSC – IT Center for Science, Finland, for computational resources. We acknowledge the support and the use of resources of Instruct-ERIC (PID6308), part of the European Strategy Forum on Research Infrastructures (ESFRI), and the Research Foundation - Flanders (FWO) for the Nanobody discovery. We acknowledge Nele Buys, Eva Beke, Alison Lundqvist and Katleen Willibal for the technical assistance during the Nanobody discovery and Nanobody workshop.

## Author Contributions

C.B., J.T.H. and W.W.Y. conceived of the study. J.T.H., K.K., J.J. and W.W.Y. devised experimental strategies. V.O.P., J.T.H, W.W.Y. supervised the project. D.T., V.O.P. and J.J.K. performed genome editing. E.P., J.J.K. and J.S. carried out nanobody generation and characterisation. H.K. carried out LC-MS/MS analysis. J.J.K., G.C., and J.T.H. performed cryo-EM and model building. J.J.K., J.T.H., W.W.Y. wrote the manuscript with contributions and critical feedback from all the authors.

## Data availability

Datasets generated during the current study are available from the Protein Data Bank (PDB) accession codes xxxx, and Electron Microscopy Data Bank (EMDB) accession codes xxxxx. All other data generated or analysed during this study are included in this published article and its Supplementary Information.

## Additional Information

Supplementary Information is available for this paper. Correspondence and requests for materials should be addressed to.

## Extended Data

**Extended Data Fig. 1.**
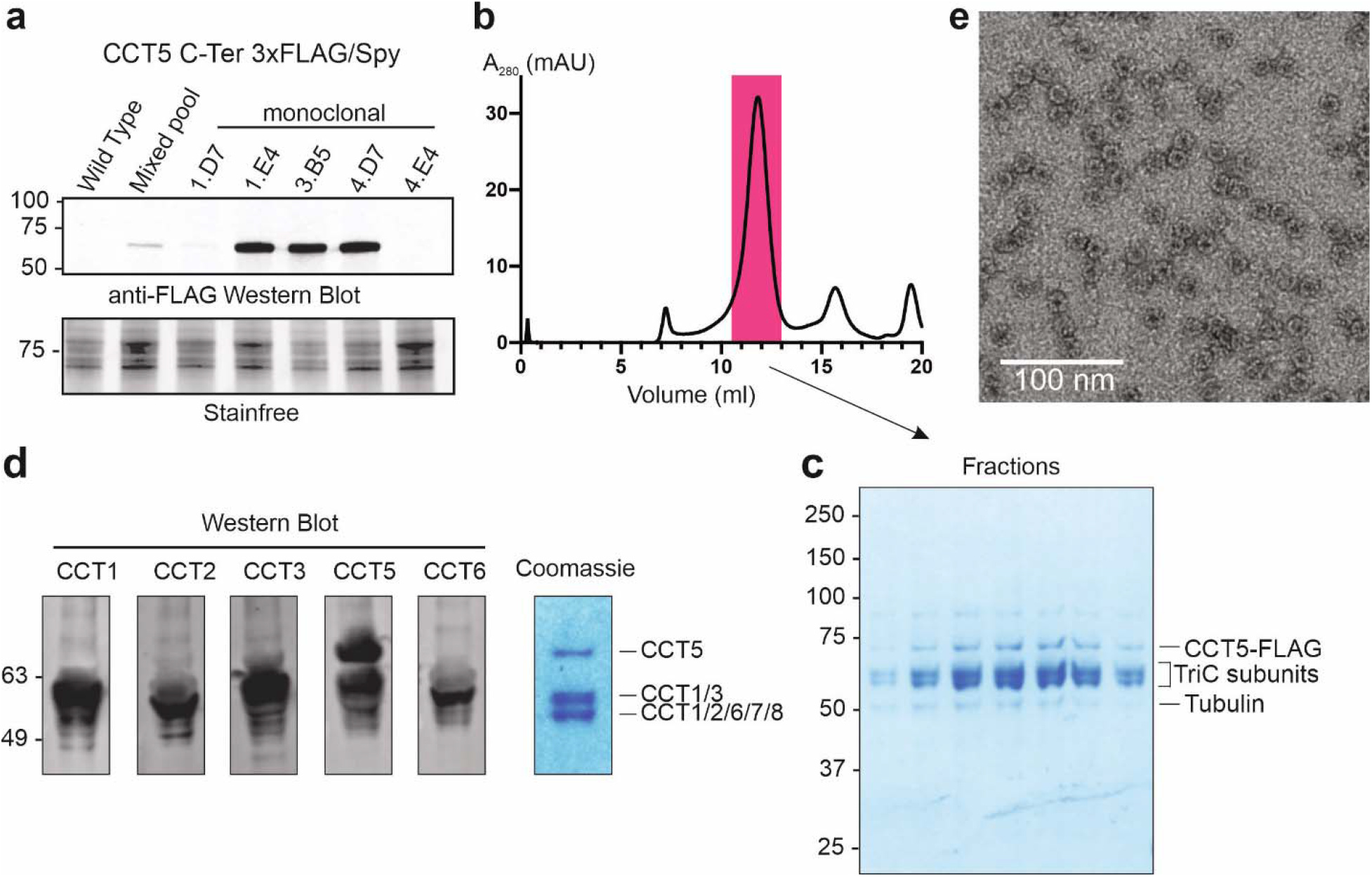
Selection and large-scale purification of monoclonal CCT5 C-term edited cell lines. (**a**) Anti-FLAG western blot (top) and stain-free imaging (bottom) of wild type HEK293T, mixed pool, and five monoclonal cell lines. (**b**) Size exclusion profile of large-scale TRiC purification with pooled fractions indicated by pink shading. (**c**) SDS-PAGE of fractions from size exclusion chromatography showing double-banding patterns of TRiC subunits, and tubulin co-elution with TRiC. (**d**) Anti-CCT western blots (left) and Coomassie visualisation (right) of monoclonal 3.B5 line showing presence of TRiC subunits. Coomassie gel bands were excised, digested with trypsin and analysed using LC-MS/MS to confirm presence of TRiC subunits. (**e**) Negative stain micrograph showing well-defined TRiC particles.

**Extended Data Fig. 2.**
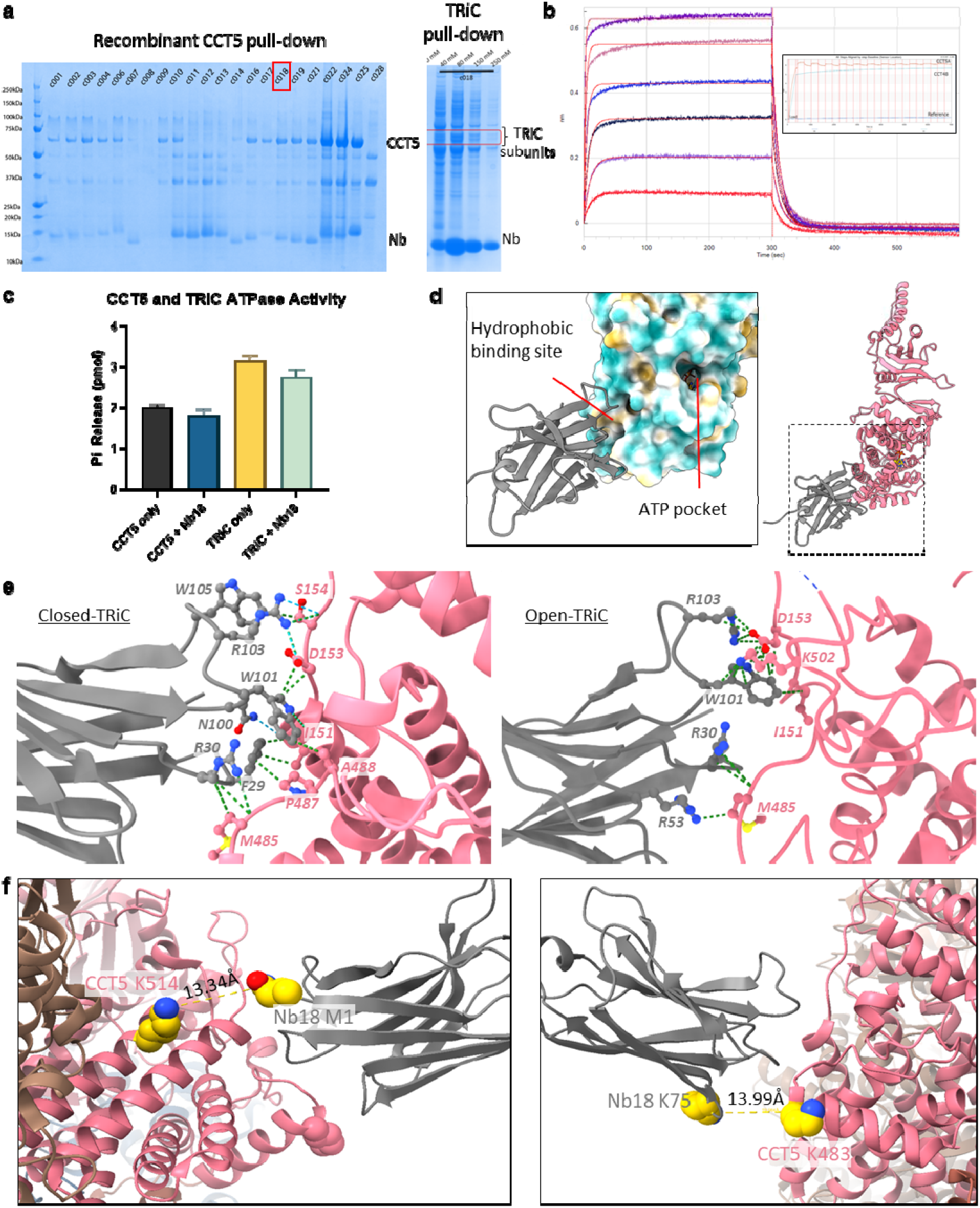
Nanobody Nb18 characterisation. (**a**) Nb18 (red box) pulls down over-expressed CCT5 (*left*) and endogenous TRiC (*right*) from affinity chromatography. (**b**) Nb18 binds recombinant CCT5 in biolayer interferometry (Kd = 86 nM). *Inset*: Raw sensogram showing that recombinant CCT4 does not bind Nb18. (**c**) CCT5 and TRiC ATPase activity in the presence of Nb18. Nb18 was added at 5:8 molar ratio to recombinant CCT5, and at 1:1 to endogenous TRiC (n=3 distinct samples with SD). (**d**) Nb18 binds to a hydrophobic interface of CCT5 in proximity to ATP binding pocket. (**e**) Interacting residues between Nb18 and CCT5 in closed-TRiC (*left*) and open-TRiC (*right*). Green dashed lines indicate van der Waals contacts and blue dashed lines indicate hydrogen bonds. (**f**) Structural representation of CCT5-Nb18 crosslinks. Crosslinked residues are highlighted as spheres in gold. Dotted lines indicate distances between amines.

**Extended Data Fig. 3.**
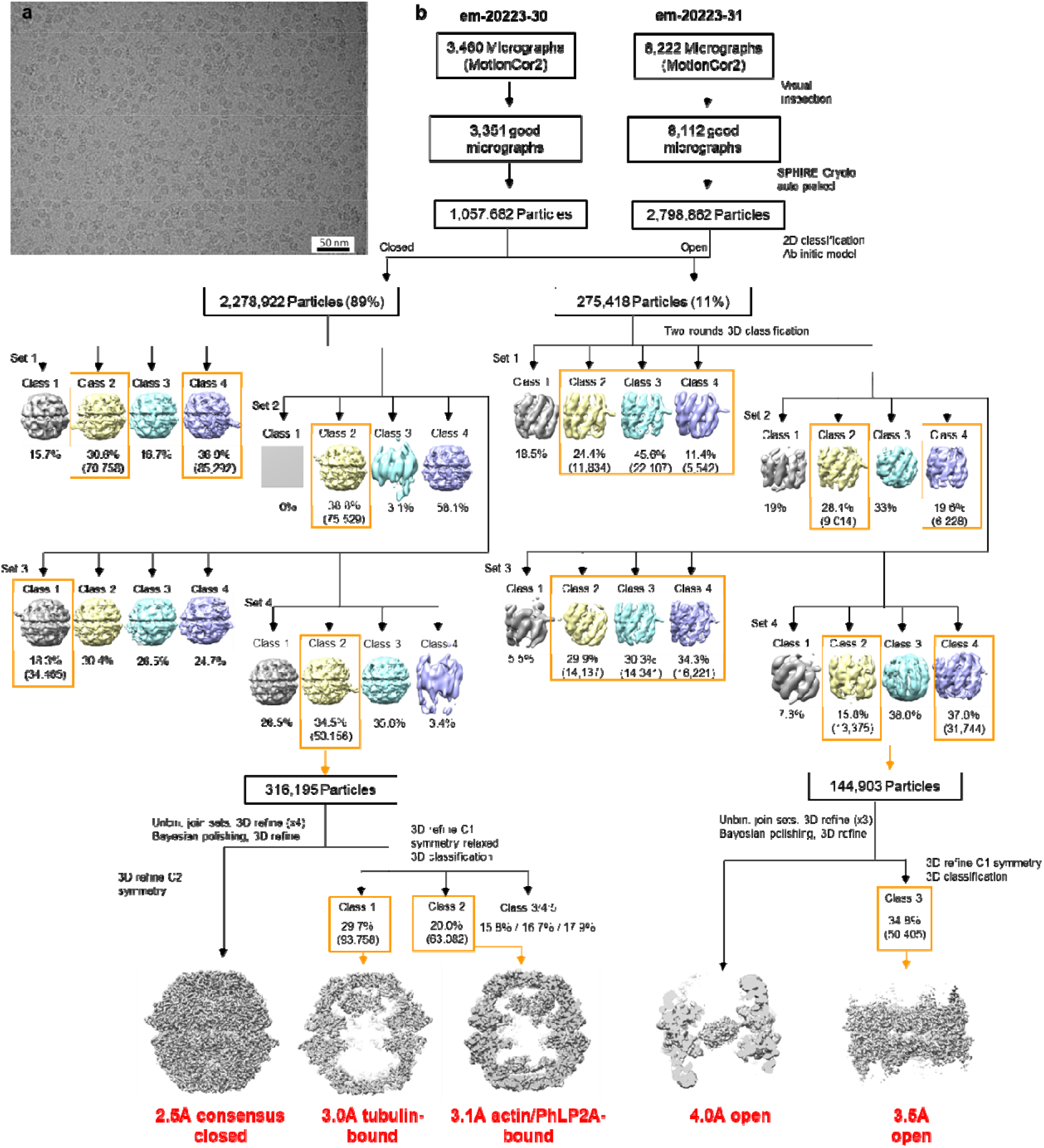
EM data processing. (**a**) A representative micrograph of purified TRiC complexes. (**b**) Cryo-EM data processing workflow. During the processing steps, the data were split in four sets (Set 1-4) which were processed in parallel for computational efficiency.

**Extended Data Fig. 4.**
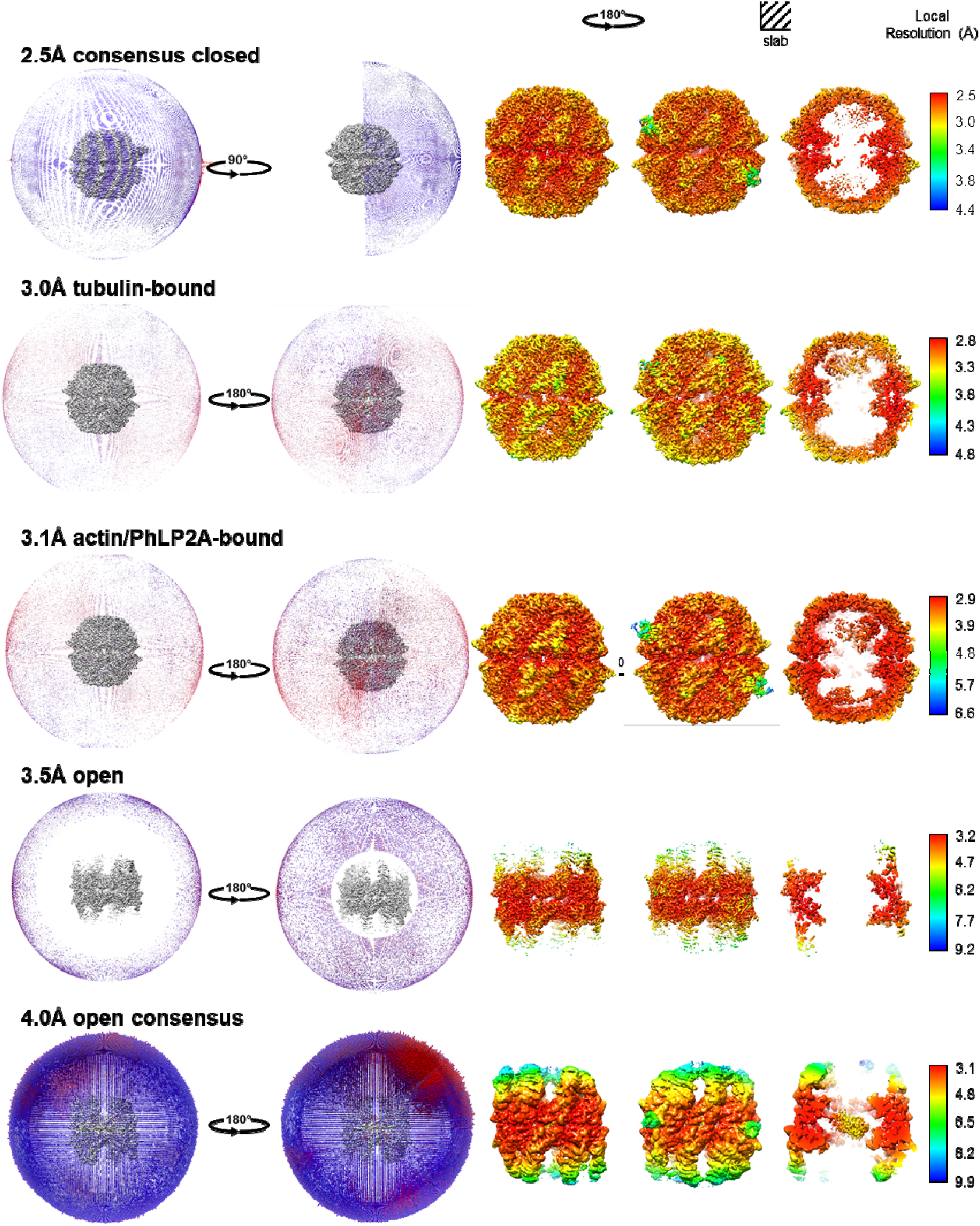
Map resolutions. Angular distribution and local resolution of all maps

**Extended Data Fig. 5.**
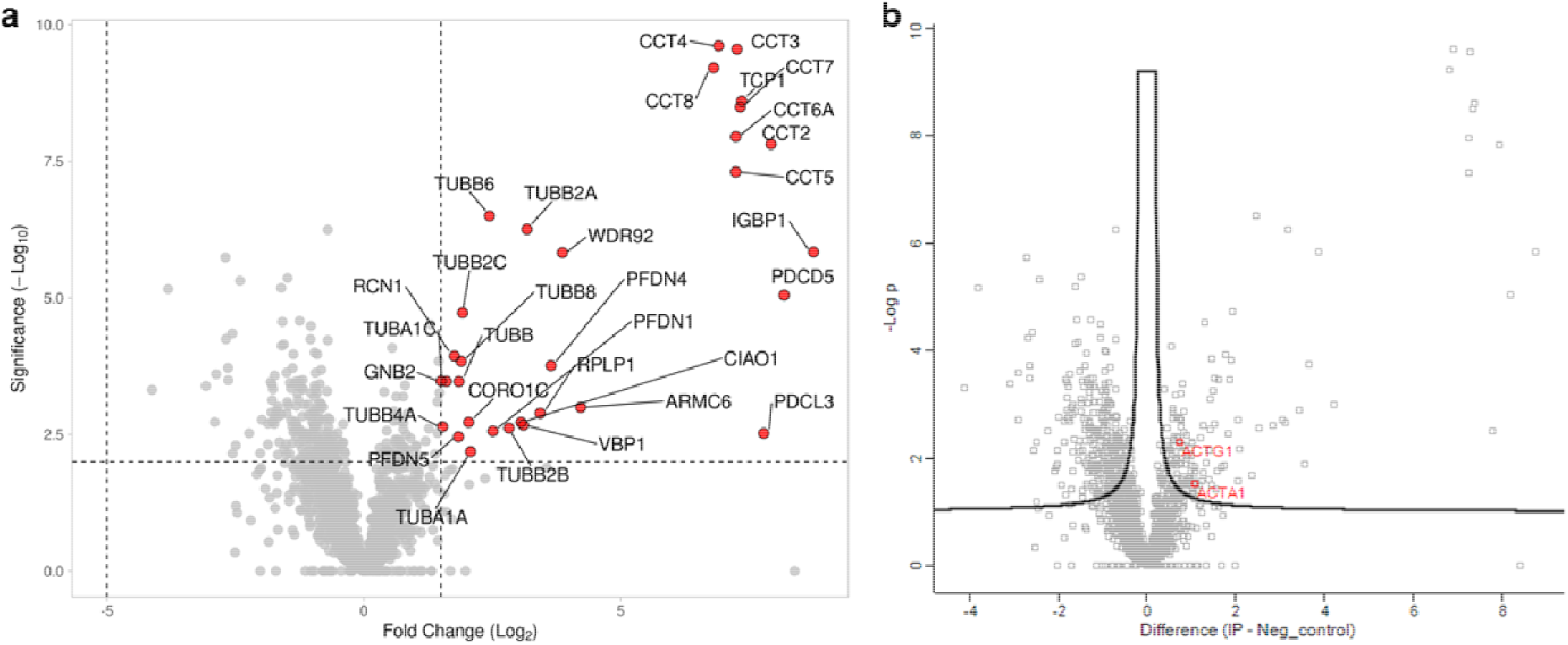
Volcano plot of peptides identified in solution digest LC-MS/MS. (**a**) Peptides are plotted according to intensity significance vs fold change over control sample. Fold change threshold set to >1.5 and significance threshold set to 2. (**b**) Peptides are plotted according to intensity significance vs fold change over control sample. Black line denotes peptides that are significantly enriched/reduced with FDR = 0.05 and s0 = 0.1. According to log2 difference, actin was slightly enriched in two peptide identifications: one that could not distinguish between α-skeletal muscle ACTA1, α-cardiac muscle 1 ACTC1, aortic smooth muscle ACTA2, γ-enteric smooth muscle ACTG2, and cytoplasmic 2 isoforms ACTB (shown in the figure as ACTA1), and one that identified cytoplasmic 2 actin (ACTG1). Raw data are found in Supplementary Data 1.

**Extended Data Fig. 6.**
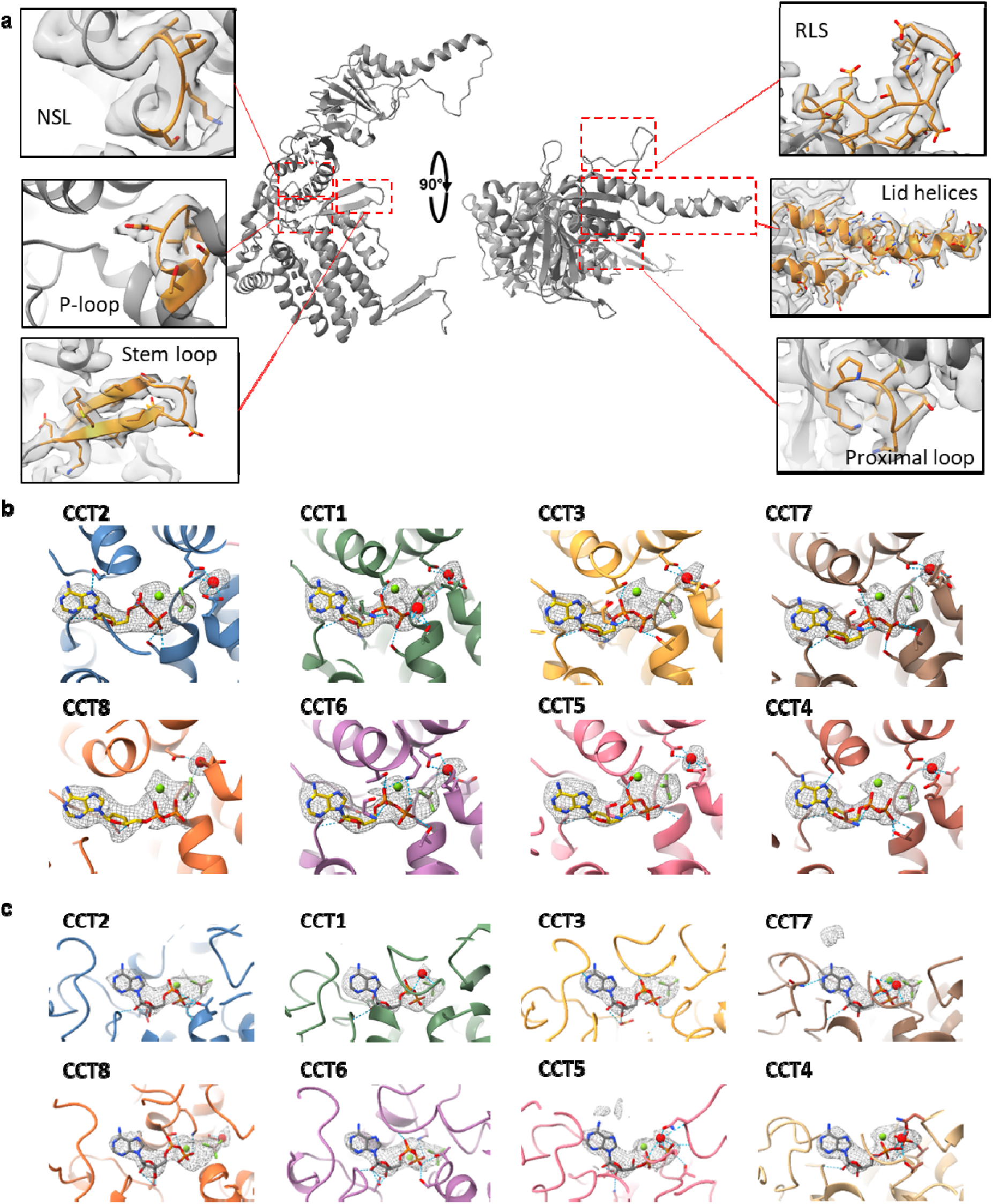
Conserved structural features of TRiC. (**a**) Conserved structural motifs common to all CCT subunits (here represented by CCT1) including apical lid helices, proximal loop, release loop of substrate (RLS), P-loop, nucleotide-sensing loop (NSL), and stem loop. *Insets*: Motifs are coloured orange and density map is shown grey. (**b,c**) Snapshots of ligand density in the nucleotide binding site for each CCT subunit of (b) closed-TRiC and (c) open-TRiC. Dashed lines represent hydrogen bonding. ADP molecules are shown in yellow with Mg^2+^ in green and water in red.

**Extended Data Fig. 7.**
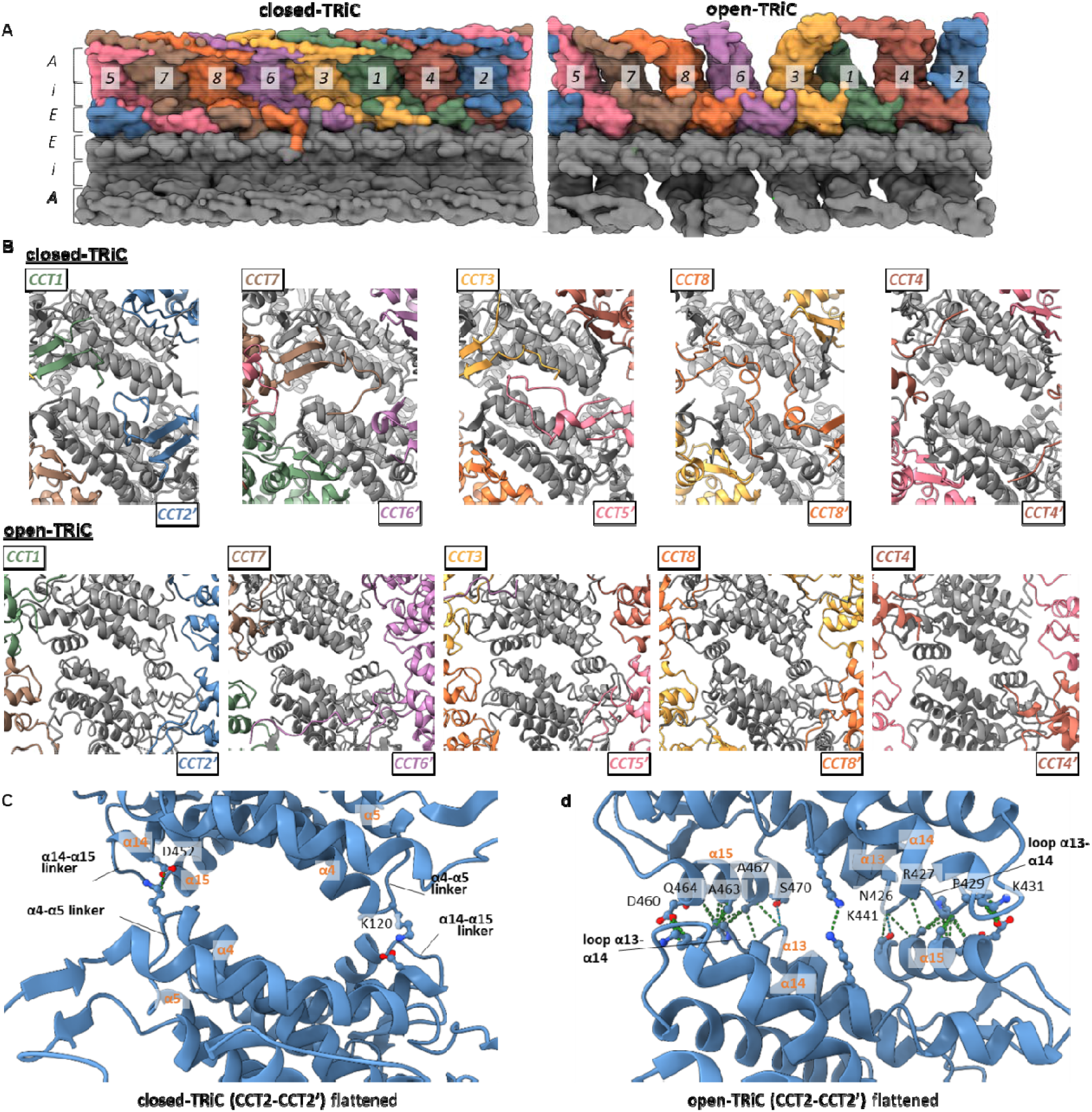
Intra- and inter-ring subunit contacts. (**a**) Flattened surface representation showing conformational differences between closed-TRiC and open-TRiC in the equatorial (*E*), intermediate (*I*) and apical (*A*) domains. (**b**) Views of the network of N-terminal extensions between subunits across the ring interface in closed-TRiC state (*top*). Note that these interactions were not observed in open-TRiC state (*bottom*) due to disorder of the N-terminal extensions. (**c,d**) Flat rendering of the CCT2-CCT2’ inter-ring stacking in closed-TRiC (c) and open-TRiC (d). Compared to closed-TRiC, rearranged helix α15 pushes loop α4-α5 from the *trans* subunit away from the interface. Inter-ring contacts are now mediated between the entire helical face of α15, and loop α13-α14 from the *trans* subunit.

**Extended Data Fig. 8.**
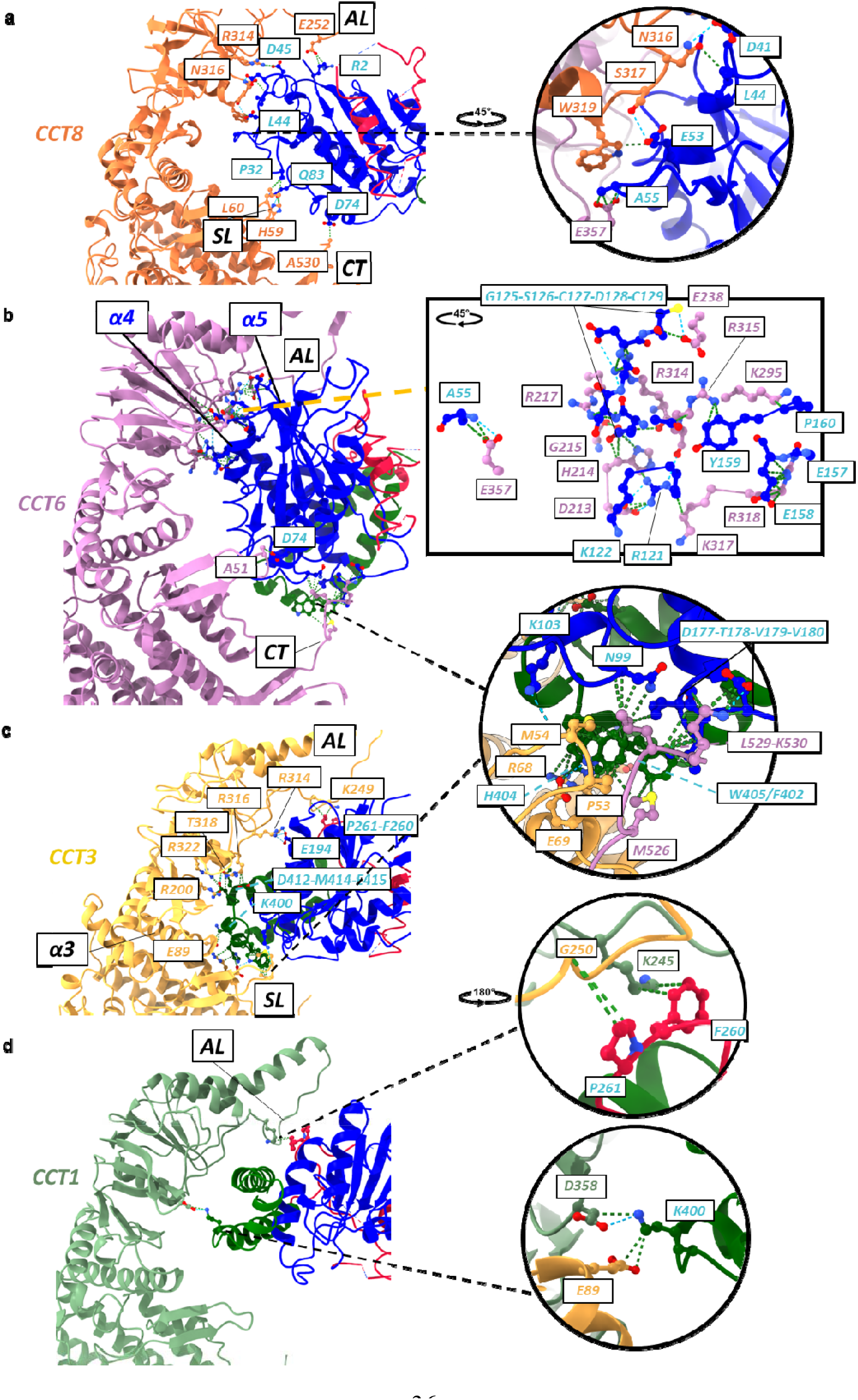
Tubulin-TRiC interactions. (**a**) Side view of CCT8 (red) binding to tubulin. Tubulin is coloured by domain including N-term domain (blue), TBD (crimson), and C-term domain (dark green). Residues are labelled by CCT colour or in cyan to represent tubulin. Inset (rotated 45°) shows network of interactions between CCT8 and CCT6 (purple). Green dashes indicate residue contacts; blue dashes indicate hydrogen bonds. Structural motifs are labelled as apical loop (AL), stemloop (SL), C-terminus (CT) and helix *X* (α*X*). (**b**) Side view of CCT6 binding to tubulin. Inset shows large interaction network between tubulin N-term domain and CCT6 apical domain displayed as atoms. (**c**) Side view of CCT3 (yellow) binding to tubulin. Inset shows interaction between C-terminus of CCT6, stem loop of CCT3, and tubulin regions ranging from the N-term domain and C-term domain. (**d**) Side view of CCT1 (green) binding to tubulin. Inset (top; rotated 180°) shows interactions between TBD and C-term tubulin domains, CCT3 apical domain, and CCT1 apical domain. Inset (bottom) shows interactions between C-term tubulin domain with intermediate domains of CCT1 and CCT3.

**Extended Data Fig. 9.**
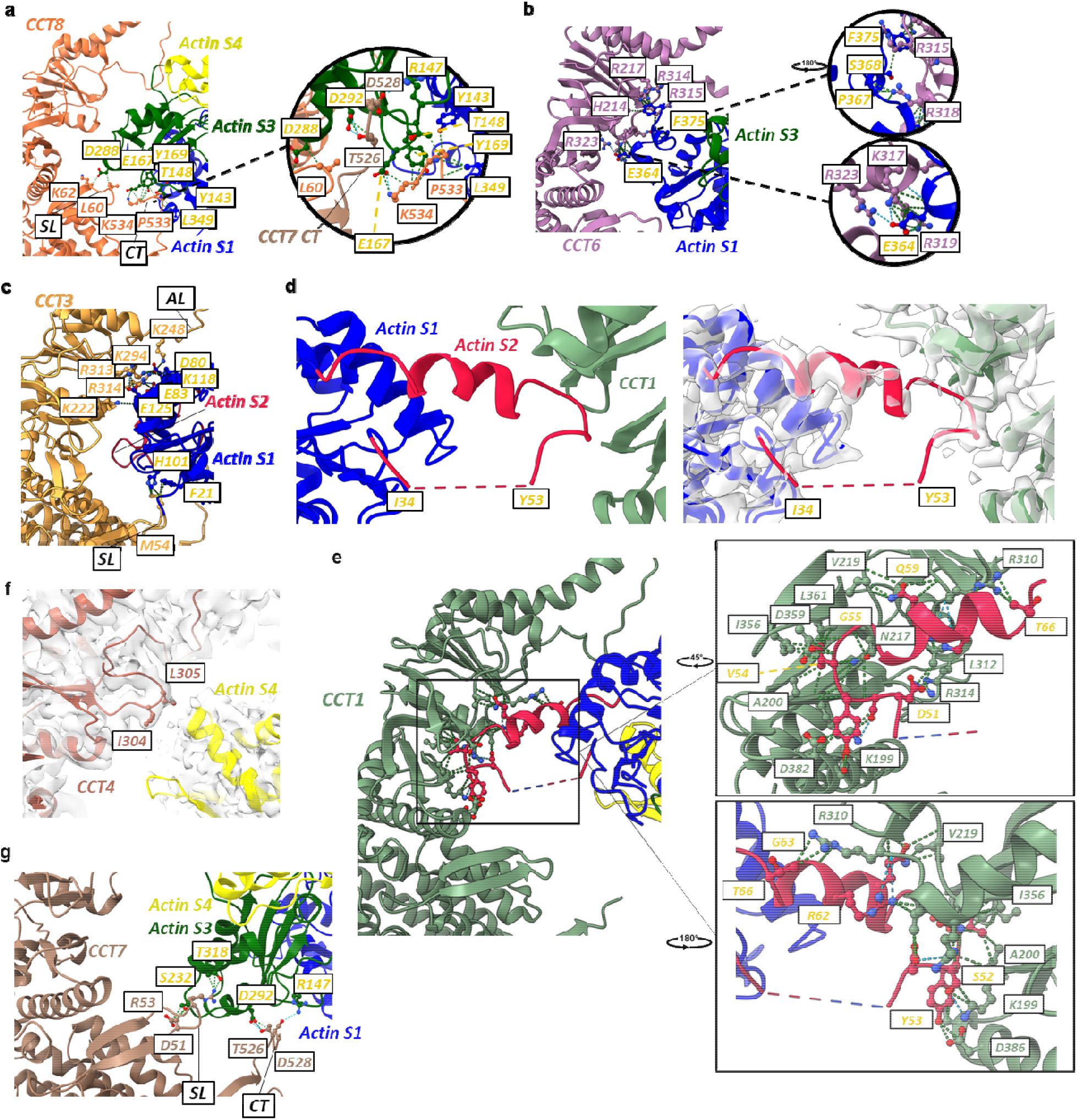
Actin-TRiC interactions. (**a**) Side view of CCT8 (orange) binding to actin. (**b**) Side view of CCT6 (purple) binding to actin. Inset (top; rotated 180°) shows zoomed in interactions near C-terminus of actin. Inset (bottom) shows residue contacts between actin residue Glu364 and CCT6. (**c**) Side view of CCT3 (yellow) binding to actin. (**d**) Zoomed in view of subdomain 2 binding within groove formed by CCT1 (green) and missing density between residues 34-53 without (*left*) and with map (*right*). (**e**) Side view of CCT1 (green) binding to actin. Insets (top rotated 45°; bottom rotated 180°) show actin subdomain 2 helix binding within groove of CCT1. (**f**) Side view of CCT4 (dark red) binding to actin. (**g**) Side view of CCT7 (brown) binding to actin. *For all panels*: actin is coloured by domain including subdomain 1 (S1, blue), subdomain 2 (S2, crimson), subdomain 3 (S3, dark green), and subdomain 4 (S4, yellow); residues are labelled by CCT colour or in gold to represent actin; green dashes indicate van der Waals contacts; blue dashes indicate hydrogen bonds; structural motifs are labelled as apical domain loop (AL), stem-loop (SL) and C-terminus (CT).

**Extended Data Fig. 10.**
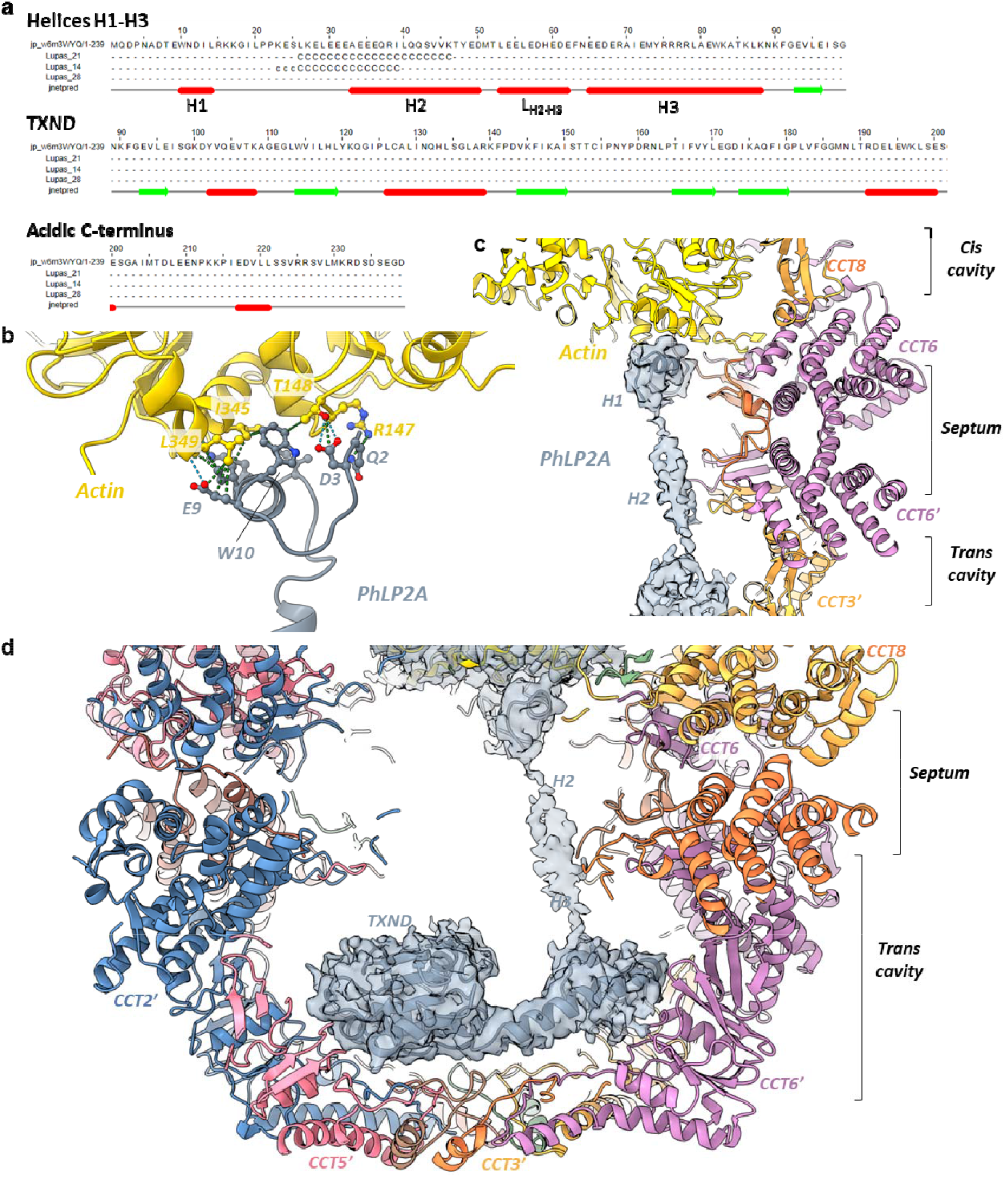
Structure of PhLP2A within TRiC chamber. (**a**) Secondary structure prediction of human PhLP2A. The three N-terminal helices H1-H3 are labelled. (**b**) Interactions between actin and PhLP2A helix H1. (**c**) Side slice of TRiC central chamber, showing the spread of PhLP2A (shown as grey density) across both *cis*- and *trans*-cavities. (**d**) Side slice of the TRiC *trans*-cavity occupied by helix H3 and TXND of PhLP2A.

## Supplementary Information

### Supplementary Tables

**Supplementary Table 1.** Cryo-EM data collection, refinement and validation statistics.

|  | Closed,<br>consensus<br>(EMDB-xxxx)<br>(PDB xxxx) | Closed,<br>tubulin-bound<br>(EMDB-xxxx)<br>(PDB xxxx) | Closed,<br>Actin/PhLP2A-bound<br>(EMDB-xxxx)<br>(PDB xxxx) | Open<br>(EMDB-xxxx)<br>(PDB xxxx) |
| --- | --- | --- | --- | --- |
| **Data collection and processing** |  |  |  |  |
| Magnification | 81,000 | 81,000 | 81,000 | 81,000 |
| Voltage (kV) | 300 | 300 | 300 | 300 |
| Electron exposure (e-/Å<sup>2</sup>) | 62 | 62 | 62 | 62 |
| Defocus range (μm) | 0.75 to 2.5 | 0.75 to 2.5 | 0.75 to 2.5 | 0.75 to 2.5 |
| Pixel size (Å) | 1.06 | 1.06 | 1.06 | 1.06 |
| Symmetry imposed | C2 | C1 | C1 | C1 |
| Initial particle images (no.) | 3,856,544 | 3,856,544 | 3,856,544 | 3,856,544 |
| Final particle images (no.) | 316,195 | 93,758 | 63,082 | 144,903 |
| Map resolution (Å) | 2.5 | 3.0 | 3.1 | 3.5 |
| FSC threshold | 0.143 | 0.143 | 0.143 | 0.143 |
| Map resolution range (Å) | 2.5–4.4 | 2.8–4.8 | 2.9–6.6 | 3.1–9.9 |
| Map sharpening *B* factor (Å<sup>2</sup>) | -63.0 | -65.3 | -57.7 | -70.2 |
| **Refinement** |  |  |  |  |
| Initial model used (PDB code) | 6KS6 | 6KS6 | 6KS6 | 6NRA |
| Model-to-map resolution (Å) | 2.64 | 3.05 | 3.27 | 3.51 |
| FSC threshold | 0.5 | 0.5 | 0.5 | 0.5 |
| Model-to-map correlation<br>(Phenix) | 0.73 | 0.86 | 0.74 | 0.81 |
| Model resolution range (Å) |  |  |  |  |
| Model composition |  |  |  |  |
| Non-hydrogen atoms | 66,477 | 69,139 | 71,009 | 34,246 |
| Protein residues | 8626 | 8970 | 9219 | 4480 |
| Ligands | 48 | 48 | 48 | 44 |
| *B* factors (Å<sup>2</sup>) |  |  |  |  |
| Protein | 72.62 | 72.55 | 70.77 | 64.35 |
| Ligand | 171.45 | 171.45 | 158.28 | 136.15 |
| R.m.s. deviations |  |  |  |  |
| Bond lengths (Å) | 0.007 | 0.004 | 0.008 | 0.005 |
| Bond angles (°) | 0.827 | 0.607 | 0.863 | 0.816 |
| Validation |  |  |  |  |
| MolProbity score | 1.64 | 1.77 | 1.85 | 2.07 |
| Clash score | 4.63 | 6.80 | 7.32 | 6.81 |
| Poor rotamers (%) | 1.73 | 1.75 | 1.55 | 1.87 |
| Ramachandran plot |  |  |  |  |
| Favored (%) | 96.57 | 96.69 | 95.6 | 92.01 |
| Allowed (%) | 3.33 | 3.22 | 4.29 | 7.81 |
| Disallowed (%) | 0.09 | 0.09 | 0.11 | 0.18 |

**Supplementary Table 2.** List of inter-subunit cross links (white shade) and TRiC-Nb18 cross links (grey shade) identified by XL-MS.

| Protein 1 | Site 1 | Protein 2 | Site 2 |
| --- | --- | --- | --- |
| CCT1 | K126 | CCT4 | K55 |
| CCT6 | K199 | CCT8 | K400 |
| Nb18 | M1 | CCT5 | K514 |
| Nb18 | K75 | CCT5 | K483 |

### Supplementary Data

**Supplementary Data 1 List of peptides identified in LC-MS/MS.** Table includes columns denoting significance of data, −Log (p-value), difference value based on intensity difference between TRiC and control samples (positive values = enriched, negative values = downregulation), protein IDs and protein names, gene names, and details on number of peptides, sequence coverage, and intensity.

### Supplementary Notes

**Supplementary Note 1: Analysis of LC-MS/MS data** To validate substrates bound within our cryo-EM map densities, we prepared two sets of samples for solution digest LC-MS/MS including one set from the CCT5-FLAG cell line and one set from wild-type HEK293T cells. Four biological replicates of each were affinity purified using anti-FLAG affinity gel matrix and elutions were digested with trypsin and analysed by LC-MS/MS. In the LC-MS/MS data (Supplementary Data 1) it was observed that tubulin and other CCT-associated proteins were enriched in the CCT5-FLAG samples as compared to wild-type samples. In addition, several peptides were present in all four samples isolated from the CCT5-FLAG cell line, while not being observed in negative control samples isolated from WT HEK293T cells. These peptides are derived from proteins including: PCNA interacting partner (PARPBP), Target of rapamycin complex subunit LST8 (MLST8), Estradiol 17-beta-dehydrogenase 8 (HSD17B8), Actin-related protein 2/3 complex subunit 1B (ARPC1B), Cullin-associated NEDD8-dissociated protein 1 (CAND1), Cyclic AMP-dependent transcription factor ATF-7 (ATF7), NEDD8 (NEDD8), Denticleless protein homolog (DTL), Tubulin-folding cofactor B (TBCB), Regulator of MON1-CCZ1 complex (C18orf8), and Phosducin-like protein (PDCL3).

**Supplementary Note 2: Analysis of TRiC-tubulin interactions** In general, the tubulin-CCT contacts are formed with the subunit apical and intermediate domains, with a few contributed from the stem-loops and C-termini. Tubulin contacts several subunits, to varying degree, as described below using the *TUBB2A* sequence. Going around the TRiC ring in a clockwise manner viewed in top-down orientation, tubulin localizes to CCT8, CCT6, CCT3, and CCT1 (Fig. 2d).

CCT8 interacts with tubulin modestly using the stem-loop (His59 and Leu60 of CCT8 with Pro32 and Glu83 of tubulin), intermediate domain (Arg314 and Arg316 of CCT8 with Asp45 and Leu44 of tubulin, respectively), and there is one contact between the apical domain helical protrusion loop (Glu252 of CCT8) and the N-terminus of tubulin (Arg2; Extended Data Fig. 8a). Of note, there is an interaction network between the intermediate domain of CCT8 (Asn316-Ser317-Trp319), the apical domain of CCT6 (Glu357), and an N-terminal domain loop of tubulin (Glu53-Ala55; Extended Data Fig. 8a, inset).

CCT6 participates the most in tubulin binding and the main interaction site is a negatively charged region of the beta-sheets of the CCT6 apical domain that sits just beneath the helical protrusion of the apical domain with helices α4 and α5 of tubulin (Extended Data Fig. 8b, inset). Two equatorial regions also interact with tubulin, including the stem-loop of CCT6 (Ala51) with Asp74 of tubulin, and, interestingly, the C-terminus of CCT6 (Extended Data Fig. 8b). The C-terminus of CCT6 is fully ordered in the tubulin-bound map up to Lys530, but can only be modelled up to residue Met526 in the consensus closed map, suggesting that the C-terminal extensions likely play a part in substrate binding as opposed to the role in inter-ring stacking seen for the N-terminal extensions. This stabilized C-terminal region of CCT6 interacts with N- and C-terminal tubulin domains including Met526 of CCT6 with Phe402/Trp405 of tubulin, Leu529 of CCT6 with Asn99//Val179/Val180/Trp405 of tubulin, and Lys530 of CCT6 with Asp177/Thr178 of tubulin, along with regions from CCT3 (Extended Data Fig. 8c, inset).

Tubulin contacts with CCT3 include the stem-loop (Pro53/Met54) with Trp405 of tubulin, and helix α3 of CCT3 (Arg68) with His404 of tubulin. Other interacting regions between CCT3 and tubulin include the CCT3 equatorial domain helix α3 (Glu89) with Lys400 of tubulin, and intermediate domain of CCT3 (Arg200/Arg316/Thr318/Arg322) with Asp412/Met414/Glu415 of tubulin (Extended Data Fig. 8c). CCT3 also interacts in two locations with both tubulin and CCT1: the apical lid loops of both CCT3 (Gly250 of CCT3 with Pro261 of tubulin) and CCT1 (Lys245 of CCT3 with Phe260 of tubulin) along with helix α3 of CCT3 (Glu89 of CCT3 with Lys400 of tubulin) and the stem-loop of CCT1 (Asp358 of CCT1 with Lys400 of tubulin; Extended Data Fig. 8d, insets).

**Supplementary Note 3: Analysis of TRiC-actin interactions** Interactions between individual subunits of TRiC and actin are described below, in clockwise TRiC subunit orientation using the *ACTB* sequence for actin.

CCT8 interacts with subdomain 3 through its stem-loop (Leu60/Lys62) with Asp288 of actin (Extended Data Fig. 9a), and its C-terminus (Extended Data Fig. 9a, inset). The C-terminal binding includes the following contacts with actin: Gly531/Gly532/Pro533 of CCT8 contact Leu349 in subdomain 1 of actin, and Pro533 of CCT8 additionally contacts Tyr143 from subdomain 1 as well as Glu167 from subdomain 3. Lys534 of CCT8 also interacts with subdomain 3 including residues Glu167 and Tyr169. Interestingly, the C-terminus of CCT7 is also involved in coordinating actin in this region Extended Data Fig. 9a, inset; Extended Data Fig. 9g).

CCT6 interacts exclusively with the C-terminus of actin in subdomain 1 (Extended Data Fig. 9b), through the same interface in the intermediate domain as the tubulin-bound model (Extended Data Fig. 8b). This includes a positively charged region of CCT6 containing Arg314, Arg217, and His214 that bind to the C-terminal Phe375 residue of actin. Additionally, contacts are formed between Arg315 of CCT6 and Ser368 of actin, Arg318 of CCT6 and Gly366/Pro367 of actin (Extended Data Fig. 9b, inset top), and interactions are observed between Glu364 of actin and Lys317/Arg319/Arg323 of CCT6 (Extended Data Fig. 9b, inset bottom).

CCT3 interacts with actin subdomain 1, through its stem-loop (Met54 of CCT3 with Phe21/His101 of actin) and apical domain. The apical domain interactions consist of a similar positively charged interface to CCT6 and includes Lys222/Arg313 of CCT3 with Glu125 of actin, Arg314 of CCT3 with Lys118/Glu83 of actin, and Lys248/Lys294 of CCT3 with Asp80 of actin (Extended Data Fig. 9c).

CCT1 interacts with the main region of actin subdomain 2 at the site of the disordered D-loop that plays a role in ATP binding (Extended Data Fig. 9d). The interaction between CCT1 and actin spans residues Asp51 to Thr66 of actin (Extended Data Fig. 9e). This region interacts with a groove in the intermediate/lower apical domain of CCT1 and includes 12 CCT1 residue contacts and nine actin subdomain 2 residue contacts (Extended Data Fig. 9e, insets), suggesting CCT1 plays an important role in sequestering this disordered region during actin folding.

While interactions between CCT4 and actin were difficult to determine due to the weak density exhibited in subdomain 4 of actin, Ile304 and Leu305 of CCT4 appeared to engage the density near subdomain 4 (Extended Data Fig. 9f). This is the only region of CCT4 that could interact with actin and suggests that subdomain 4 is not well ordered and only associates loosely with CCT4 to position the actin monomer for subsequent folding. CCT7 makes several contacts with actin subdomain 3 including the stem-loop (Asp51 of CCT7 with Ser232 of actin, and Arg53 of CCT7 with Thr318 of actin) and through its C-terminus (Thr526 of CCT7 with Asp292 of actin, and Asp528 of CCT7 with Arg147 of actin; Extended Data Fig. 9g), in the same regions as that of CCT8 (Extended Data Fig. 9a). Taken together, these interactions display an actin monomer in its near-native fold that is coordinated by specific interactions with the CCT6 hemisphere of TRiC in the closed state.

